# Deep Learning Reveals Endogenous Sterols as Allosteric Modulators of the GPCR-Gα Interface

**DOI:** 10.1101/2023.02.14.528587

**Authors:** Sanjay Kumar Mohanty, Aayushi Mittal, Namra, Aakash Gaur, Subhadeep Duari, Saveena Solanki, Anmol Kumar Sharma, Sakshi Arora, Suvendu Kumar, Vishakha Gautam, Nilesh Kumar Dixit, Karthika Subramanian, Tarini Shankar Ghosh, Debarka Sengupta, Shashi Kumar Gupta, Natarajan Arul Murugan, Deepak Sharma, Gaurav Ahuja

## Abstract

Endogenous intracellular allosteric modulators of GPCRs remain largely unexplored, with limited binding and phenotype data available. This gap arises from the lack of robust computational methods for unbiased cavity identification, cavity-specific ligand design, synthesis, and validation across GPCR topology. Here, we developed Gcoupler, an AI-driven generalized computational toolkit that leverages an integrative approach combining *de novo* ligand design, statistical methods, Graph Neural Networks, and Bioactivity-based ligand prioritization for rationally predicting high-affinity ligands. Using Gcoupler, we interrogated intracellular metabolites that target and regulate the GPCR-Gα interface (Ste2p-Gpa1p), affecting pheromone-induced programmed cell death in yeast. Our computational analysis, complemented by experimental validations, including genetic screening, multi-omics, site-directed mutagenesis, biochemical assays, and physiological readouts, identified endogenous hydrophobic metabolites, notably sterols, as direct intracellular allosteric modulators of Ste2p. Molecular simulations coupled with biochemical signaling assessment in site-directed Ste2p mutants further confirmed that metabolites binding to GPCR-Gα obstruct downstream signaling, possibly via a cohesive effect. Finally, by utilizing isoproterenol-induced, GPCR-mediated human and neonatal rat cardiac hypertrophy models, we observed that elevated metabolite levels attenuate hypertrophic response, reinforcing the evolutionary relevance of this mechanism.

## INTRODUCTION

G protein-coupled receptors (GPCRs) are critical regulators of cellular processes and thus represent prime drug targets (Alhosaini et al., 2021; Calebiro et al., 2021; Yang et al., 2021). While traditional GPCR-targeted therapies focus on orthosteric sites, recent advances have revealed allosteric sites offering novel therapeutic avenues (Bourque et al., 2022; Wold and Zhou, 2018)(Bourque et al., 2022; Leach et al., 2007; Rees et al., 2002). Although exogenous synthetic allosteric modulators are known, endogenous counterparts remain poorly characterized (Doller, 2017; O’Callaghan et al., 2012; Reyes-Alcaraz et al., 2020; Stornaiuolo et al., 2015; van der Westhuizen et al., 2015; Zhang et al., 2015). Developing high-affinity endogenous modulators requires integrating structure-based design, artificial intelligence (AI), and assays, yet traditional approaches like SAR analysis are hampered by limited GPCR allosteric modulator data (Basith et al., 2018; Gupta et al., 2021). While experimental techniques like FRET and BRET can validate allosteric compounds, their use in high-throughput screening for novel intracellular modulators is challenging (Hoffmann and Bünemann, 2010; Jaeger et al., n.d.; Zhou et al., 2021). Identifying endogenous GPCR allosteric modulators is further complicated by factors like incomplete GPCR topology data (Congreve et al., 2020) and vast chemical space (Topiol, 2018a; Yang et al., 2021). This necessitates a hybrid computational approach combining allosteric site prediction, *de novo* ligand synthesis, and efficient screening, potentially enhanced by AI. Existing *de novo* drug design tools often lack practical applicability for this purpose due to computational limitations and high technical demands (Böhm, 1992; Li et al., 2021; Nishibata and Itai, 1991; Pearlman and Murcko, 1993; Sicho et al., 2021; Spiegel and Durrant, 2020; Yuan et al., 2011).

To overcome these challenges, we developed Gcoupler, a software suite (available as a Python package and Docker image) that integrates structural biology, statistical methods, and deep learning to identify GPCR allosteric modulators. We demonstrated the usability and applicability of Gcoupler in identifying novel endogenous modulators of GPCRs by exploiting the α-pheromone (α-factor)-induced mating or programmed cell death (PCD) pathway of *S. cerevisiae*. Notably, it is well documented that the highly elevated, non-physiological levels of α-factor trigger PCD in less than half of the MATa population (Zhang et al., 2006). Moreover, an equivalent concentration of α-factor triggers distinct PCD kinetics across distinct laboratory strains; for example, BY4741 is more resistant than the W303 strain (Sokolov et al., 2020). We, therefore, hypothesized that a subset of pheromone-resistant cells might regulate the Ste2p-mediated PCD signaling via the endogenous intracellular metabolites by operating at the Ste2-Gα binding interface. Using Gcoupler, we identified a subset of intracellular metabolites that could potentially bind to Ste2p (GPCR) at the Gpa1 (Gα) binding interface and obstruct the downstream signaling. Our computational results further suggest that hydrophobic ligands such as sterols strengthen the Ste2p-Gpa1p binding and might trigger a cohesive response that potentially obstructs downstream signaling. Experimental evidence further supported these findings that the elevated intracellular levels of these metabolites rescue the pheromone-induced PCD. To evaluate the evolutionary conservation and possible clinically relevant translation of this mechanism, we tested these metabolites on human and rat isoproterenol-induced, GPCR-mediated cardiac hypertrophy model systems and observed attenuated response in the cardiomyocytes pretreated with GPCR-Gα-protein interface modulating metabolites.

## RESULTS

### Development and validation of Gcoupler

Designing novel target molecules by integrating the topological, chemical, and physical attributes of protein cavities necessitates advanced neural networks. While existing approaches like Bicyclic Generative Adversarial Networks (BicycleGANs) (Skalic et al., 2019) and Recurrent Neural Networks (RNNs) (Xu et al., 2021) have demonstrated potential, end-to-end standalone tools for GPCR-specific ligand design remain scarce. To address this, we developed the Gcoupler and provided it to the community as a Python Package and a Docker image. Gcoupler adopts an integrative approach utilizing structure-based, cavity-dependent *de novo* ligand design, robust statistical methods, and highly powerful Graph Neural Networks. Gcoupler consists of four interconnected modules, i.e., Synthesizer, Authenticator, Generator, and BioRanker, that collectively impart a smoother, user-friendly, and minimalistic experience for the end-to-end *de novo* ligand design.

Synthesizer, the first module of Gcoupler, takes a protein structure as input in Protein Data Bank (PDB) format and identifies putative cavities across the protein surface, providing users with the flexibility to select cavities based on druggability scores or user-supplied critical residues. Since cavity-dependent molecule generation mainly depends on the chemical composition and geometric constraints of the cavity, it is, therefore, indispensable to select the cavity for the downstream steps considering its chemical nature (hydrophobicity/hydrophilicity) and functional relevance (proximity to the active site or residue composition), among others. Accounting for these, Gcoupler offers flexibility to the users to select either of its predicted cavities based on the user-supplied critical residue or by user-supplied cavity information (amino acids) using third-party software (e.g., Pocketome) (Hedderich et al., 2022). To enhance user experience, Gcoupler computes and outputs all identified cavities along with their druggability scores using LigBuilder’s V3 (Yuan et al., 2020) cavity module. Briefly, these druggability scores consider solvent accessibility, cavity exposure or burial, and detected pharmacophores and cavities, which are further prioritized based on this score. Post cavity selection, the Synthesizer module generates cavity-specific ligands influenced by topology and pharmacophores, outputting SMILES, cavity coordinates, and other requisite files to downstream modules for further steps **(Figure 1a)**. The chemical composition of the *in silico* synthesized ligands by the Synthesizer module is influenced by the cavity topology (3D) and its composition (pharmacophores). Noteworthy, the Synthesizer module of Gcoupler employs LigBuilder V3 (Yuan et al., 2020), which utilizes the genetic algorithm for the *in silico* ligand synthesis. Notably, the fragment library of LigBuilder, comprising 177 distinct molecular fragments in Mol2 format, allows the selection of multiple seed structures and extensions that best complement the cavity pharmacophores throughout multiple iterative runs. For each run, once a seed structure is confirmed, Gcoupler employs a hybrid approach of the Growing and Linking modes of the LigBuilder build module, enabling the stepwise addition of small fragments to the seed structure within the binding pocket of the target GPCR to build synthetic ligands. Gcoupler generates 500 unique molecules by default, though it can also be user-defined. The Synthesizer module of Gcoupler enhances LigBuilder V3 practical applicability through automation, dynamic adaptability, and abstraction. This allows for more efficient and targeted ligand generation, even in challenging design scenarios for GPCR ligand design. However, it lacks user-defined library screening, proposes synthetically challenging molecules, and often requires post-processing to isolate high-affinity binders from a broad affinity range of synthetically designed compounds.

**Figure 1:**
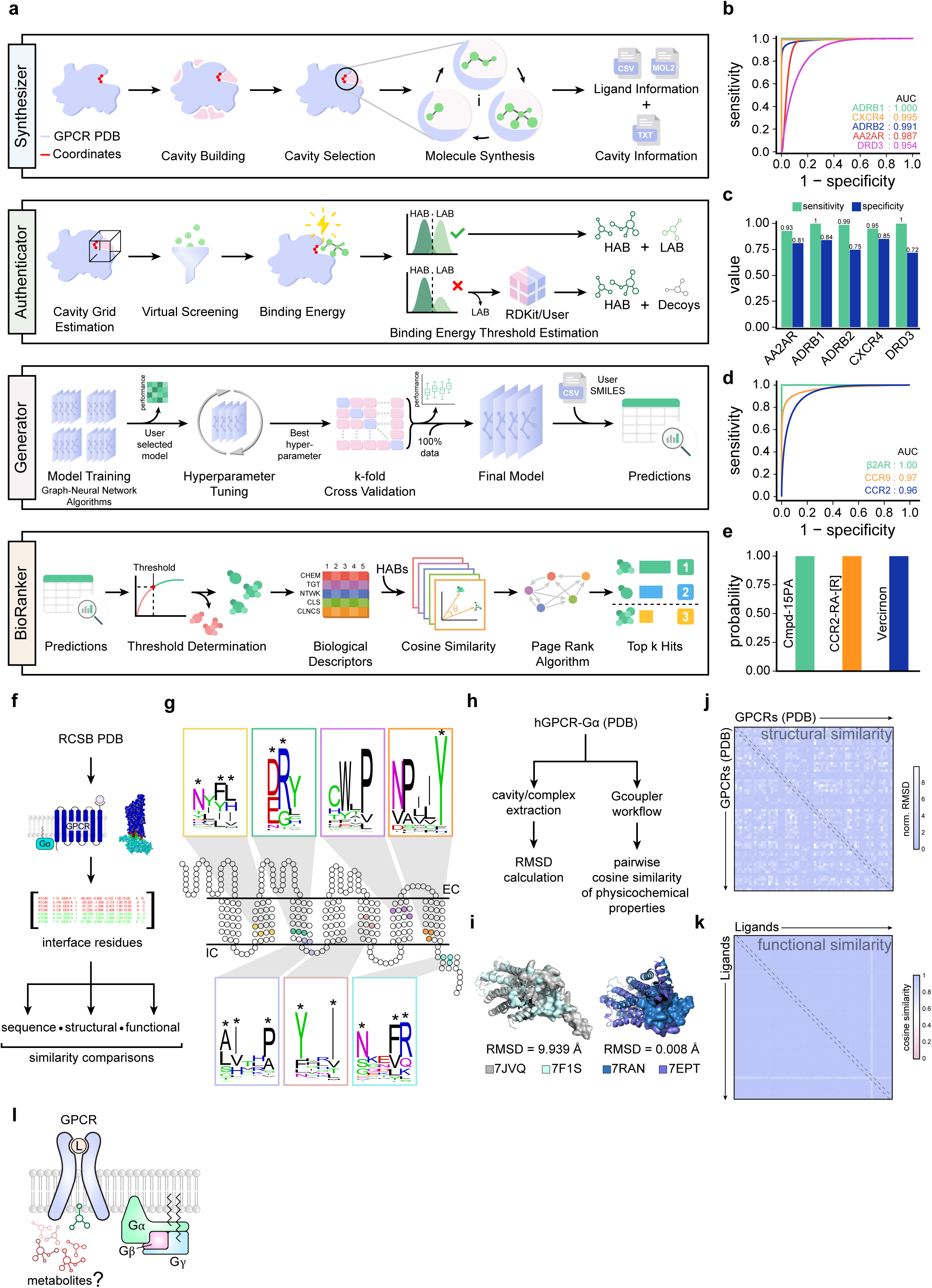
Development, Benchmarking and Validation of Gcoupler Computational Framework. **(a)** Schematic workflow depicting different modules of the Gcoupler package. Of note, Gcoupler possesses four major modules, i.e., Synthesizer, Authenticator, Generator and BioRanker. **(b)** AUC-ROC curves of the finally selected model for each of the indicated GPCRs. Note: Experimentally validated active ligands and decoys were used in the testing dataset. **(c)** Bar graphs depicting the sensitivities and specificities of the indicated GPCRs with experimentally validated active ligands and reported decoys. **(d)** The AUC-ROC curve indicates the model’s performance in the indicated conditions. **(e)** Bar graphs indicating the prediction probabilities for each experimentally validated ligand. **(f)** Schematic workflow illustrates the steps in measuring and comparing the structural conservation of the GPCR-Gα-protein interfaces across human GPCRs. **(g)** Snakeplot depicting the standard human GPCR two-dimensional sequence level information. Conserved motifs of the GPCR-Gα-protein interfaces are depicted as WebLogo. Asterisks represent residues of conserved motifs present in the GPCRs-Gα-protein interfaces. Of note, the location of the motifs indicated in the exemplary GPCR snake plot is approximated. **(h)** Schematic workflow illustrates the steps in measuring and comparing the structural conservation of the GPCR-Gα-protein interfaces across human GPCRs. **(i)** Representative structures of the proteins depicting highly conserved (low RMSD) and highly divergent (high RMSD) GPCR-Gα-protein interfaces. PDB accession numbers are indicated at the bottom. **(j)** Heatmap depicting the RMSD values obtained by comparing all the GPCR-Gα-protein interfaces of the available human GPCRs from the protein databank. Of note, the RMSD of the Gα-protein cavity was normalized with the RMSDs of the respective whole proteins across all pairwise comparisons. **(k)** Heatmap depicting the pairwise cosine similarities between the *in silico* synthesized ligands of the GPCR-Gα-protein interfaces of the available human GPCRs using Gcoupler. **(l)** Schematic diagram depicting the hypothesis that the intracellular metabolites could allosterically modulate the GPCR-Gα interaction.

To address this limitation, we developed the second module of Gcoupler, termed Authenticator. This module processes output files from the Synthesizer module, conducting downstream validation steps and preparing results for constructing Deep Learning-based classification models (third module). The Authenticator requires input protein 3D structure in PDB format, cavity coordinates, and all silico-generated molecules from the synthesizer module. Authenticator utilizes this information to further segregate the synthesized compounds into high-affinity binders (HAB) and low-affinity binders (LAB) by leveraging a structure-based virtual screening approach (AutoDock Vina) (Trott and Olson, 2010) and statistically-backed hypothesis testing for distribution comparisons **(Figure 1a)**. The Authenticator module outputs the free binding energies of all the generated compounds, which further segregates the compounds into HABs and LABs by the statistical submodule while ensuring the optimal binding energy threshold and class balance. Of note, the Authenticator is also capable of leveraging the Empirical Cumulative Distribution Function (ECDF) for binding energy distribution comparisons of HABs and LABs and performs the Kolmogorov–Smirnov test (Berger and Zhou, 2014), Epps-Singleton test (Goerg and Kaiser, 2009), and Anderson-Darling Test (Engmann and Cousineau, 2011) for hypothesis testing. This expanded array of statistical tests allows users to employ methodologies that best suit their data distribution characteristics, ensuring robust and comprehensive analyses. Moreover, the Authenticator module incorporates a unique feature for decoy synthesis using high-affinity binders. This functionality enables the generation of a negative dataset in scenarios where the Synthesizer module fails to produce an optimal number of LABs. By synthesizing decoys from high-affinity binders, users can effectively balance their datasets, enhancing the reliability of downstream analyses. Lastly, the Authenticator module also accommodates user-supplied negative datasets as an alternative to LABs (Mysinger et al., 2012). This feature provides users with the flexibility to incorporate external data sources, enabling robust prediction model building by the subsequent Generator module.

The Generator, the third module, employs state-of-the-art GNN models such as Graph Convolution Model (GCM), Graph Convolution Network (GCN), Attentive FP (AFP), and Graph Attention Network (GAT) to construct predictive classifiers using Authenticator-informed classes. These GNN algorithms are tailored to extract features from the graph structure of the compounds generated by the Synthesizer and apply them to the classification task by leveraging Authenticator-informed class information. For instance, the Graph Convolution Model assimilates features by analyzing neighboring nodes, while the Graph Convolution Network extracts features through a convolutional process. The Attentive FP model focuses attention on specific graph segments, and the Graph Attention Network employs attention mechanisms to learn node representations. By default, Generator tests all four models using standard hyperparameters provided by the DeepChem framework (https://deepchem.io/), offering a baseline performance comparison across architectures. This includes pre-defined choices for node features, edge attributes, message-passing layers, pooling strategies, activation functions, and dropout values, ensuring reproducibility and consistency. All models are trained with binary cross-entropy loss and support default settings for early stopping, learning rate, and batch standardization where applicable. Gcoupler provides off-the-shelf hyperparameter tuning to ensure adequate training, which is essential for optimizing model performance. After selecting the best parameters and classification algorithm, Gcoupler further ensures the mitigation of overfitting and provides a more precise estimate of model performance through k-fold cross-validation. Notably, by default, Gcoupler employs three-fold cross-validation, but users can adjust this parameter.

Finally, BioRanker, the last module, prioritizes ligands through statistical and bioactivity-based tools. The first level ranking offered by BioRanker is composed of a statistical tool that encompasses two distinct algorithms, namely G-means and Youden’s J statistics, to assist users in identifying the optimal probability threshold, thereby refining the selection of high-confidence hit compounds (**Supplementary Figure 1a**). Additionally, bioactivity embeddings computed via Signaturizer (Bertoni et al., 2021) enable multi-activity-based ranking using a modified PageRank algorithm. Briefly, the bioactivity descriptors of the predicted compounds are projected onto various biological activity spaces, including Chemistry, Targets, Networks, Cells, and Clinics, by performing pairwise cosine similarity comparisons with High-Affinity Binders (HABs). The PageRank algorithm is then applied for activity-specific ranking and supports multi-activity-based ranking for sequential screening based on user-defined biological properties. BioRanker also offers flexibility through customizable probability thresholds, enabling stringent or relaxed selection of compounds. Users can also input SMILES representations for direct screening, bypassing prediction probabilities. Taken together, Gcoupler is a versatile platform supporting user-defined inputs, third-party tools for cavity selection, and customizable statistical analyses, enhancing its adaptability for diverse ligand design and screening tasks. This integrated framework streamlines cavity-specific ligand design, screening, and ranking, providing a comprehensive solution for GPCR-targeted drug discovery.

To evaluate Gcoupler’s performance, we tested its modules across five GPCRs (AA2AR, ADRB1, ADRB2, CXCR4, and DRD3) using experimentally validated ligands and matched decoys from the DUD-E dataset (Mysinger et al., 2012). The DUD-E datasets contain five GPCRs alongside information about their cavity coordinates, positive ligands, and decoys (https://dude.docking.org/subsets/gpcr). We used these five GPCRs as independent samples to evaluate different modules and sub-modules of Gcoupler. We first checked whether the cavity search algorithm of Synthesizer could accurately detect a given orthosteric ligand-binding site for a GPCR. Gcoupler accurately identified orthosteric ligand-binding sites and additional allosteric cavities across all targets, validating its *de novo* cavity detection algorithm **(Supplementary Figure 1b)**. We next asked whether Gcoupler could also synthesize molecules similar to the reported ligands for respective orthosteric sites based on the cavity’s physical, chemical, and geometric properties. For orthosteric sites, the Synthesizer module generated ∼500 compounds per GPCR. Subsequently, as per the Gcoupler default workflow, the Authenticator module conducted a virtual screening of these newly synthesized compounds, segregating them into high-affinity binders (HAB) and low-affinity binders (LAB). Although the Authenticator module provides flexibility in selecting an optimal threshold to distinguish HAB and LAB, we chose the default cut-off of −7 kcal/mol for AA2AR, CXCR4, and DRD3. For ADRB1 and ADRB2, we selected a threshold of −8 kcal/mol to minimize overlap in distributions and thus avoid class imbalance, a critical parameter that could influence the downstream model generation using the Generator module **(Supplementary Figure 1c)**. Statistical validation confirmed significant separation between these groups (p < 0.0001), enabling the Generator module to construct graph-based classification models with high values of AUC-ROC (>0.95), sensitivity, and specificity **(Figure 1b-c, Supplementary Figure d-e)**. These models reliably distinguished ligands from decoys, demonstrating Gcoupler’s accuracy in identifying high-affinity ligands.

In addition to evaluating Gcoupler’s performance for the orthosteric sites of GPCRs, we also validated its capability to identify allosteric sites and their corresponding ligands. In this case, we first gathered information about the experimentally validated GPCR-ligand complexes sourced from the PDB database. We chose three GPCR-ligand complexes (β2AR-Cmpd-15PA, CCR2-CCR2-RA-[R], CCR9-Vercirnon) from the PDB (Shen et al., 2023). We removed the ligands from the PDB files and executed the standard Gcoupler workflow with default parameters. Gcoupler successfully identified allosteric binding sites and generated classification models for synthetic compounds with consistently high AUC-ROC values (>0.95) **(Figure 1d, Supplementary Figure 2a-c)**. This high level of accuracy indicates the robustness of Gcoupler’s algorithms in distinguishing between true positives (allosteric ligands) and true negatives (non-binders). Projection of experimentally validated ligands onto these models further confirmed their predictive accuracy **(Figure 1e)**, underscoring Gcoupler’s robustness and versatility for orthosteric and allosteric ligand discovery.

Next, to evaluate the efficiency of Gcoupler, we compared its run time with the biophysics-based gold standard molecular docking (AutoDock) (Morris et al., 2009). To address the runtime efficiency, we first utilized the ChEMBL31 database (Gaulton et al., 2012) to identify GPCRs with the highest number of reported experimentally validated agonists. We selected the alpha-1A adrenergic receptor (ADRA1A) since it qualifies for this criterion and contains 993 agonists **(Supplementary Figure 3a-b)**. Methodologically, we followed the conventional steps of AutoDock Tools for molecular docking while keeping track of execution time for each step throughout the entire process until completion **(Supplementary Figure 3a, c).** In parallel, we applied the same timestamp procedure for Gcoupler, including its individual module sub-functions **(Supplementary Figure 3a-d)**. Gcoupler was 13.5 times faster, leveraging its deep learning-based Generator module and AutoDock Vina’s efficiency. Both methods provided comparable predictions for active compounds, demonstrating Gcoupler’s speed and accuracy, making it ideal for large-scale ligand design and drug discovery **(Supplementary Figure 3e-h)**.

Finally, we used Gcoupler to evaluate the ligand space conservation (functional conservation) of the GPCR-Gα interface. Specifically, we aimed to explore the possibility of direct small molecule binding to the GPCR-Gα interface to modulate downstream signaling pathways. We analyzed multiple human GPCR-Gα complexes from the PDB **(Figure 1f, Supplementary Table 1)**, identified conserved motifs (DRY, CWxL, NPxxY) and binding pockets through sequence and structural analyses **(Figure 1g)**. To determine the topological similarity of the GPCR-Gα protein interface, we undertook a detailed structural analysis across a wide array of GPCR-Gα protein complexes. This analysis involved identifying and extracting the cavities present within each complex. By focusing on these critical regions, we aimed to assess the degree of structural conservation and quantify it through normalized Root Mean Square Deviation (RMSD) values. Specifically, the normalized RMSD values, which provide a measure of the average distance between atoms of superimposed proteins, indicated a high degree of similarity. The mean RMSD value was found to be 1.47 Å, while the median RMSD value was even lower at 0.86 Å. These values suggest that the overall topology of the GPCR-Gα interface is well conserved across different complexes, highlighting the robustness of this interaction site. **(Figure 1h-j, Supplementary Figure 3i-k, Supplementary Table 2)**. Finally, to test whether this topological and sequence conservation also impacts the ligand profiles that could potentially bind to this interface, we performed the Gcoupler workflow on all 66 GPCRs and synthesized ∼50 unique ligands per GPCR **(Figure 1h).** We next computed and compared the physicochemical properties (calculated using Mordred (Moriwaki et al., 2018)) of these synthesized ligands and observed high cosine similarity, which further supports the functional conservation of the GPCR-Gα interface **(Figure 1h, k, Supplementary Figure 3l, Supplementary Table 3)**. In summary, we used Gcoupler to systematically evaluate and analyze the ligand profiles of the GPCR-Gα-protein interface and observed a higher degree of sequence, topological, and functional conservation.

### Gcoupler reveals endogenous, intracellular Ste2p allosteric modulators

We next utilized Gcoupler to test the hypothesis that the intracellular metabolites could potentially and directly regulate the GPCR signaling by directly interacting with the GPCR-Gα-protein interaction interface **(Figure 1l)**. To test this hypothesis, we utilized a well-characterized yeast mating pathway mediated via the Ste2p-Gpa1p interface **(Supplementary Figure 4a)**. We used Gcoupler to screen for such metabolites against the Yeast Metabolome Database (YMDB) (Jewison et al., 2012). We utilized the recently elucidated cryo-EM structure of the Ste2 protein (Velazhahan et al., 2021) and performed a small-scale molecular dynamics simulation by using a yeast phospholipid composition-based lipid bilayer environment (Kaneko et al., 1976), and subjected the simulated stable structure to the Gcoupler workflow **(Supplementary Figure 4b-i)**. This led to the identification of 17 potential surface cavities on Ste2p. Careful interrogation of the structurally supported Ste2p-Gpa1p interface revealed two distinct predicted cavities (annotated as IC4 (Intracellular Cavity 4) and IC5 (Intracellular Cavity 5)), collectively capturing >95% of the interface regions **(Figure 2a-b, Supplementary Figure 4j-n)**. Conservation analysis of these cavities across 14 yeast species further confirmed their structural significance **(Supplementary Figure 4o)**. We next synthesized ∼500 *in silico* synthetic compounds for both IC4 and IC5, each by leveraging the Synthesizer module of Gcoupler. To test whether the chemical space of these *in silico* synthesized ligands is cavity-specific, we performed a stringent evaluation by comparing the chemical heterogeneity of the 100 randomly selected synthesized ligands, each from the pool of 500 for IC4 and IC5, with 100 *de novo* synthesized ligands for an extracellular cavity (EC1). Of note, EC1 does not harbor any overlapping residue with IC4 or IC5 and possesses distinct pharmacophore properties **(Supplementary Figure 5a, d-e)**. Next, we computed the atom pair fingerprints and visualized the chemical heterogeneity in the low-dimensional space using 2D and 3D PCA **(Supplementary Figure 5b-c)**. These results suggest that the Synthesizer module of Gcoupler generated cavity-specific ligands by leveraging both the cavity topology (3D) and its composition (pharmacophore). We further assessed the reproducibility of the Gcoupler by synthesizing 100 in silico compounds per run across five runs for the Ste2p IC4 cavity. Visualization of the chemical heterogeneity between the compounds generated via different runs in the low-dimensional space using 2D/3D PCA and pairwise Tanimoto Similarity using atom pair fingerprints suggests heterogeneous and overlapping chemical composition among the synthesized ligands across all five runs **(Supplementary Figure 5f-h)**.

**Figure 2:**
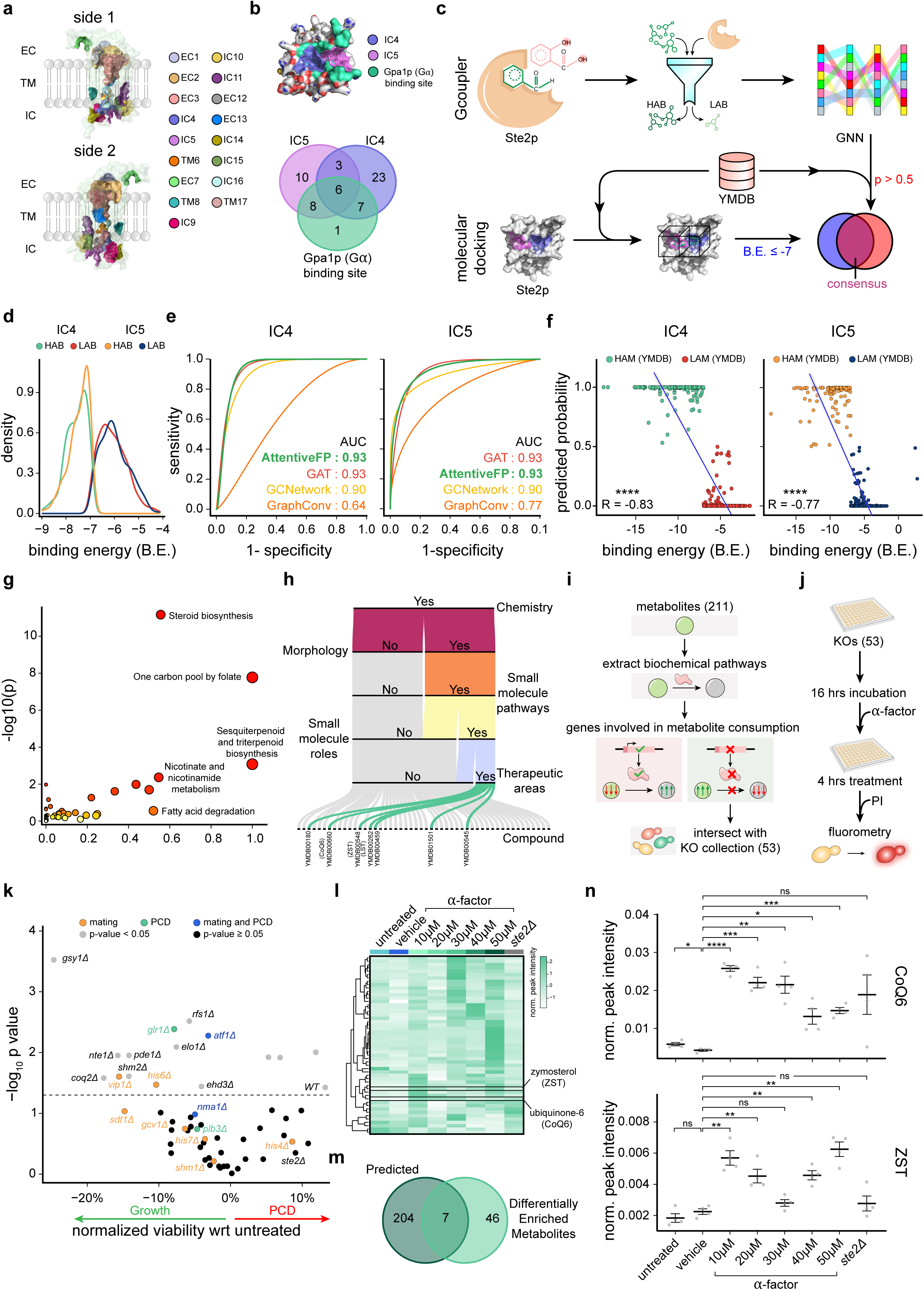
Identification of Endogenous, Intracellular Allosteric Modulators of Ste2p Using Gcoupler. **(a)** Schematic diagram depicting the topology of all the cavities identified using the Synthesizer module of the Gcoupler Python package. Of note, the cavity nomenclature includes the cavity location, i.e., EC (extracellular), IC (intracellular), and TM (transmembrane), succeeded by a numerical number. **(b)** Diagram depicting the three-dimensional view of the Ste2 protein, with highlighted Gα-protein binding site (Gpa1) and the Gcoupler intracellular cavities (IC4 and IC5). The Venn diagram at the bottom depicts the percentage overlap at the amino acid levels between the Gα-binding site and predicted IC4 and IC5. **(c)** Schematic representation of the overall workflow used to predict the endogenous intracellular allosteric modulators of Ste2 receptor using Gcoupler and molecular docking technique. Of note, Yeast Metabolome DataBase (YMDB) metabolites were used as query compounds. **(d)** Overlapping density plots depicting and comparing the distributions of synthetic compounds predicted to target the IC4 and IC5 of the Ste2 receptor using the Gcoupler package. Of note, the Authenticator module of Gcoupler segregated the synthesized compound for each cavity (IC4 or IC5) into High-Affinity Binders (HAB) and Low-Affinity Binders (LAB). **(e)** AUC (Area under the curve) plots representing the performance of the indicated models. Notably, the models were trained using the cavity-specific synthetic compounds generated using the Gcoupler package. **(f)** Scatter plots depicting the relationship (correlation) between the binding prediction probabilities using Gcoupler and binding free energies computed using molecular docking (AutoDock). **(g)** Scatterplot depicting the Pathway Over Representation Analysis (ORA) results of the endogenous metabolites that were predicted to bind to the GPCR-Gα-protein (Ste2p-Gpa1p) interface using both Gcoupler and molecular docking. **(h)** Alluvial plot showing five-level sub-activity spaces screening of the selected metabolites for IC4. **(i)** Schematic diagram depicting the workflow opted to narrow down on the single metabolic gene mutants. **(j)** Schematic diagram depicting the experimental design used to screen single metabolic gene mutants for α-factor-induced PCD. Cell viability was assessed using a Propidium iodide-based fluorometric assay. **(k)** Scatter plot depicting the impact of α-factor stimuli on cellular viability, assessed using Propidium iodide-based fluorometric assay. The y-axis represents −log10(p-value) of the one-sample Student’s t-test between the normalized PI fluorescence of untreated and treated conditions. The x-axis represents the percentage inhibition or increase in cellular viability, estimated using a Propidium Iodide-based assay. The mutants reported to be involved in mating, PCD, or both are indicated in orange, green, and blue, respectively. The statistically non-significant mutants are indicated below the dashed line in black. **(l)** Heatmap depicting the relative enrichment/de-enrichment of differentially enriched metabolites in the indicated conditions. Of note, four biological replicates per condition were used in the untargeted metabolomics. **(m)** Venn diagram depicting the overlap between the predicted endogenous intracellular allosteric modulators of Ste2p and differentially enriched metabolites (DEMs) identified using untargeted metabolomics. **(n)** Mean-whisker plot depicting the relative abundance of ubiquinone 6 (CoQ6) and zymosterol (ZST) in the indicated conditions. Student’s t-test was used to compute statistical significance. Asterisks indicate statistical significance, whereas ns represents non-significance.

Post these performance/reproducibility checks of Gcoupler, we segregated the 500 *in silico* synthesized ligands for IC4 and IC5, each into high-affinity binders (HABs) and low-affinity binders (LABs) by the Authenticator module of Gcoupler. Notably, the estimated binding energy threshold was set at −7 kcal/mol, a widely accepted cutoff in virtual screening (Alanzi et al., 2024; Wong et al., 2022) **(Figure 2c-d)**. A deep investigation into these classified synthetic compounds showed a comparable similarity between the HABs and LABs of the aforementioned target cavities, respectively **(Supplementary Figure 6a-c)**. The Generator module built classification models by implementing four distinct Graph Neural Network algorithms. Comparing the model performance metrics suggests that Attentive FP outperformed other algorithms for both cavities **(Figure 2e, Supplementary Figure 6d)**. Next, we screened yeast metabolites, compiled from YMDB, on the best-performing model (hyperparameter-tuned Attentive FP model) and predicted metabolites that could potentially bind to the IC4 and IC5 of the Ste2p-Gpa1p interface with a binding probability cutoff of > 0.5 **(Figure 2c)**. To further optimize the lead metabolite list, we parallelly performed the standard molecular docking using AutoDock with YMDB metabolites against IC4 and IC5 of Ste2p, respectively, and ultimately selected the consensus metabolites (binding energy ≤ −7 kcal/mol and binding probability > 0.5) for the downstream analysis **(Figure 2c, Supplementary Figure 6e, Supplementary Table 4)**. Of note, the consensus metabolites list was further segregated into HAM (High-Affinity Metabolites) and LAM (Low-Affinity Metabolites) based on the binding prediction probability (cutoff = 0.5). Comparative analysis of the binding prediction probabilities of Gcoupler and binding energies from Autodock of HAM and LAM revealed a significant negative correlation that further validates the authenticity of our novel approach **(Figure 2f).** Of note, as expected, HAMs and LAMs possess distinct atomic fingerprints, as indicated by the Principal Component Analysis **(Supplementary Figure 6f)**. Furthermore, HAMs (High-Affinity Metabolites) and LAMs (Low-Affinity Metabolites) displayed distinct atomic fingerprints, with enriched functional groups, including R2NH, R3N, ROPO3, ROH, and ROR, observed in HAMs **(Supplementary Figure 6g-h)**. To gain the pathway-level information about these putative endogenous intracellular allosteric modulators of Ste2p, we performed pathway-level over-representation analysis (Chong et al., 2018) and observed the selective enrichment of metabolites involved in the steroid, sesquiterpenoid, and triterpenoid biosynthesis and one-carbon pool by folate pathways **(Figure 2g)**. BioRanker module further pinpointed sterols, including zymosterol (ZST), ubiquinone 6 (CoQ6), and lanosterol (LST), as top candidates, exhibiting high prediction probabilities (>0.99) and structural similarity to HABs **(Figure 2h, Supplementary Figure 6i-j)**. To validate Gcoupler-identified allosteric modulators, we performed control analyses of the Authenticator and Generator modules along with blind docking of YMDB metabolites with Ste2p. In the former case, we removed the class information (HAB and LAB labels) from the *in silico* compounds synthesized for the IC4 cavity of the Ste2p, which resulted in a heterogeneous pool of chemical compounds. We next randomly split the training and testing data (5 iterations) and build independent models. Our results suggest that compared to the Authenticator-guided data splitting (HAB and LAB), the random splitting resulted in poor model performances **(Supplementary Figure 6k)**, suggesting the robustness of the Authenticator module. Additionally, we also evaluated the impact of the size of the training data on the Generator model performance. To achieve this, we randomly selected 25%, 50%, 75%, and 100% of the *in silico* synthesized compounds of the Ste2p IC4 cavity and built models using default parameters. Our results revealed a significant increase in the model performance with increased training data size **(Supplementary Figure 6l-m)**. For the latter case, we performed blind docking by AutoDock for Ste2p with the YMDB metabolites and compared these results with the cavity-specific Docking via AutoDock and Gcoupler predictions. As expected, in contrast to the cavity-specific Gcoupler and AutoDock, where we observed significant segregation of the HAM and LAM at −7 kcal/mol Binding Energy (BE) cutoff and 0.5 as the probability cutoff of Gcoupler, we failed to observe any striking differences for the HAM in the case of blind docking **(Supplementary Figure 6n)**. All these rigorous control analyses and blind docking validated the reliability of Gcoupler’s predictions, confirming its robustness in identifying cavity-specific modulators.

To experimentally validate the role of Gcoupler-predicted metabolites in Ste2p signaling, we performed a genetic screen of metabolic mutants. We first mapped the predicted allosteric-modulating metabolites to biochemical pathway databases (KEGG and MetaCyc) and identified the enzymes responsible for processing these metabolites (**Figure 2i**, **Supplementary Figure 7a, Supplementary Table 5**). Among the 53 single metabolic mutants (+ *ste2Δ*) screened, only Ste2 was previously reported in KEGG pathways for altered mating response (**Supplementary Figure 7b**). Next, we performed large-scale activity screening using α-factor-induced PCD assays with Propidium Iodide (PI), assuming that metabolic gene deletions would lead to intracellular accumulation of target metabolites. Briefly, selected single metabolic mutants (MATa) were grown under optimal conditions to late log phase (16 hours). Growth profile analysis revealed varied responses relative to wild-type, with most mutants exhibiting delayed growth kinetics (**Supplementary Figure 7c**). Late log phase cells were subsequently treated with α-factor, and PCD induction was quantified using PI-based cell viability assays (**Figure 2j**). Notably, wild-type BY4741 strains showed significant PI fluorescence increase, indicating pheromone-induced PCD, and *STE2* knockout mutants (*ste2Δ*) showed no significant death as expected. Interestingly, most metabolic mutants (94.4%) resisted α-factor-induced cell death, with some displaying accelerated growth in the presence of α-factor, indicating crosstalk between central metabolism and Ste2 signaling (**Figure 2k**). These results indicate that a significant proportion of Gcoupler predicted metabolites could directly or indirectly influence the Ste2 signaling pathway, establishing a link between metabolism and Ste2 signaling.

To further investigate the metabolic pathways associated with PCD resistance, we performed high-resolution metabolomics on cells surviving α-factor treatment at varying concentrations (**Supplementary Figure 7d**). Unbiased metabolome analysis revealed differentially enriched metabolites in the surviving population (**Figure 2l, Supplementary Figure 7e-i, Supplementary Table 6**). Cross-comparison analysis identified seven metabolites overlapping between Gcoupler predictions and survivor-enriched metabolites, with ubiquinone 6 (CoQ6) and zymosterol (ZST) showing prominent enrichment across all tested concentrations (**Figure 2m-n, Supplementary Figure 7j**). Over-Representation Analysis of the differentially-enriched metabolites suggests their involvement in glyoxylate and dicarboxylate metabolism, purine metabolism, and vitamin B6 metabolism, among others. (**Supplementary Figure 7k**). Taken together, the findings from genetic screening and untargeted metabolomics hint towards the interplay between the central metabolism and Ste2 signaling, with computationally predicted metabolites like ZST and CoQ6 potentially conferring resistance to α-factor-induced PCD.

### Elevated Endogenous Metabolic levels selectively inhibit GPCR signaling

To evaluate the stability of the interactions of zymosterol (ZST), lanosterol (LST), and ubiquinone 6 (CoQ6) at the Ste2p-Gpa1p interface, we performed three independent replicates of molecular dynamics (MD) simulations (short run of 100 ns) of the Ste2p-metabolite complex for both cavities **(Figure 3a-c, Supplementary Figure 8a)**. MD simulation results suggest that the interactions between the metabolites and Ste2p at the Ste2p-Gpa1p interface are thermodynamically stable in almost all cases across both the cavities (IC4 and IC5), except for the ubiquinone 6 (CoQ6), which harbored a fluctuating Root Mean Square Deviation (RMSD) over the simulation timeframe **(Figure 3c)**. Notably, fluctuating RMSD is observed only in the case of IC5, while it is within the permissive range for IC4 **(Figure 3c, Supplementary Figure 8b)**. We further evaluated IC4 by performing a longer simulation run for 550 ns with these three Ste2p-metabolite complexes and observed stable complexes in the case of zymosterol (ZST) and lanosterol (LST) while observing fluctuations in the ligand RMSD for ubiquinone 6 (CoQ6) post 100 ns **(Supplementary Figure 8c)**, which may be a result of its greater conformational flexibility. To gain further insight into the contributing residues from the MD simulations, we performed a residue-wise decomposition analysis that provides information about the energy contributions from the different residues to total binding free energies **(Supplementary Figure 9a)**. These results suggest that IC4 and IC5 specific residues predominantly contribute to the total binding free energies. Notably, the binding free energies are obtained as an average of over 500 configurations corresponding to the last 50 ns of the MD simulations.

**Figure 3:**
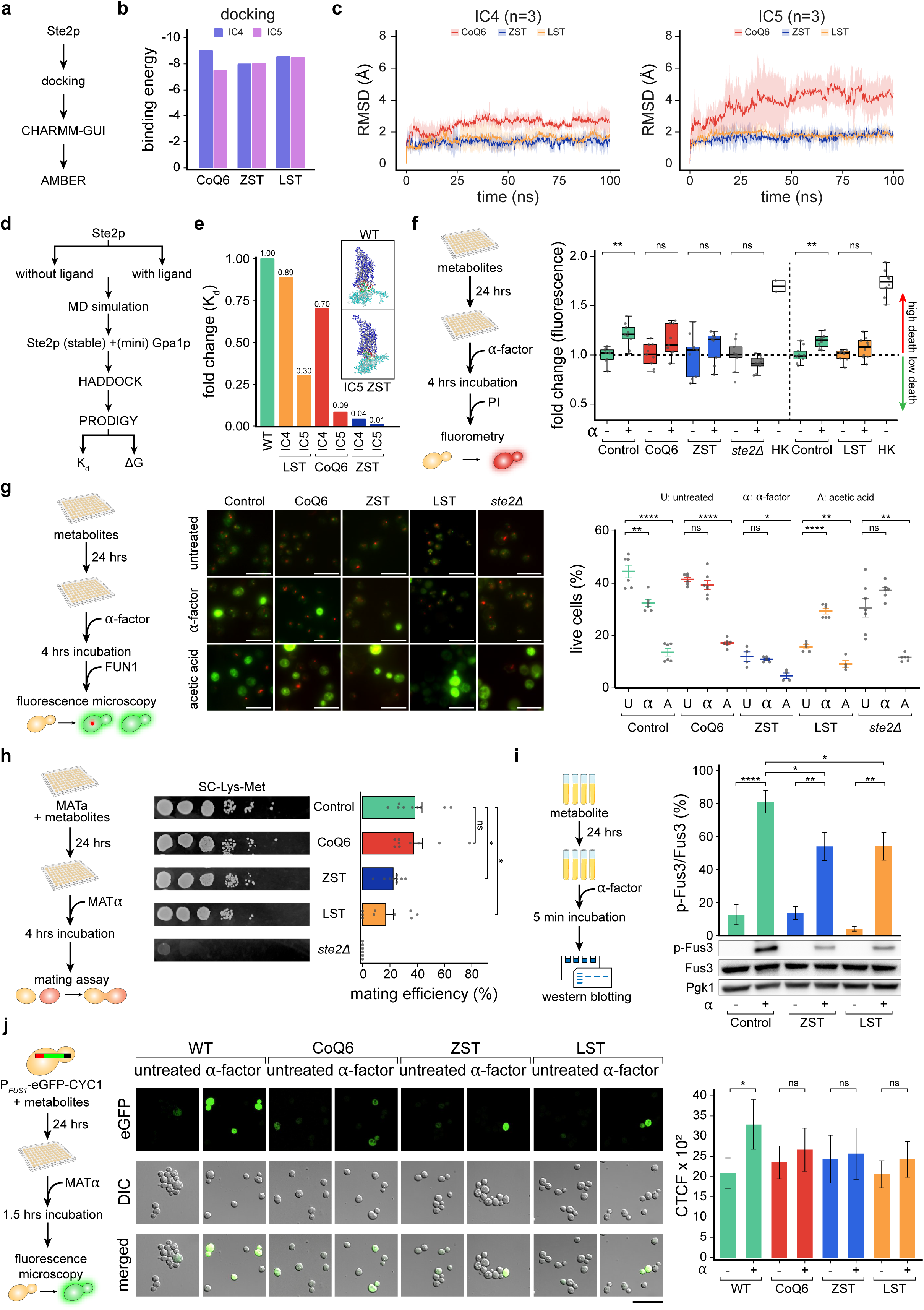
Elevated Endogenous Metabolite Levels Stabilize Ste2p-Gpa1p Interactions and Selectively Inhibit GPCR Signaling. **(a)** Scheme representing the key steps opted for preparing Ste2p structure for downstream computational analysis. **(b)** Barplots depicting the binding energies obtained by the docking of Ste2p and indicated metabolites across IC4 and IC5. **(c)** Line plots depicting the Root Mean Square Deviation (RMSD) changes over simulation timeframes from the three independent replicates of the indicated conditions in the indicated conditions. The spread of the data is indicated as Standard Deviation (SD). Notably, RMSD is provided in Angstroms (Å), whereas the simulation time is in nanoseconds (ns). **(d)** Workflow depicting the steps involved in Ste2p-miniG-protein docking using HADDOCK and PRODIGY web servers. **(e)** Barplots depicting the fold change of the dissociation constant (Kd) in the indicated conditions. Notably, fold change was computed with respect to the wild-type condition (Ste2p-miniG-protein). Inlets represent molecular representations of Ste2p-miniG-protein and the highlighted interface residues. **(f)** The schematic diagram depicts the experimental workflow used to quantify α-factor-induced PCD using a propidium iodide-based cell viability fluorometric assay. Box plot on the right depicting the rescue from the α-factor-induced PCD in the indicated conditions as inferred using propidium iodide-based cell viability fluorometric assay (n=9 or 10 biological replicates; Heat-killed = 2). The y-axis represents the fold change of the propidium iodide fluorescence values with respect to their respective controls. Mann Mann-Whitney U test was used to calculate statistical significance. Asterisks indicate statistical significance, whereas ns represents non-significance. **(g)** Schematic representation (left) of the experimental approach used to measure cell vitality and viability using microscopy-based FUN^TM^1 staining. Representative micrographs (right) depicting the FUN^TM^1 staining results in the indicated conditions, Scale 10 µm. Mean-whisker plot depicting the relative proportion of the vital and viable yeast cells observed using FUN^TM^1 staining in the indicated conditions (n = 3 biological replicates). A Student’s t-test was used to compute statistical significance. Asterisks indicate statistical significance, whereas ns represents non-significance. Error bars represent the standard error of the mean (SEM). **(h)** Schematic representation (left) of the experimental design for the mating assay (n = 3 biological replicates, each with three technical replicates). MATa yeast cells were preloaded with the metabolites and then mated with MATα cells to evaluate the mating efficiency. Representative micrographs in the middle qualitatively depict the mating efficiency in the indicated conditions. The bar plots on the right depict the mating efficiency (mean ± SEM) in the indicated conditions. Student’s t-test was used to compute statistical significance. Asterisks indicate statistical significance, whereas ns represents non-significance. **(i)** Schematic representation depicting the experimental design of phospho-MAPK activity-based Western blot. Barplots depicting the p-Fus3 levels (mean ± SEM) in the indicated conditions. Error bars represent the standard error of the mean (SEM). A Student’s t-test was used to compute statistical significance. Asterisks indicate statistical significance, whereas ns represents non-significance. **(j)** Schematic representation (left) of the experimental approach used to measure the fluorescence in P*_FUS1_*-eGFP-CYC1 yeast cells. Representative micrographs (right) depicting the eGFP expression in the yeast cells in the indicated conditions. Scale 20 µm. Barplot depicting the Corrected Total Cell Fluorescence (CTCF) value (mean ± SEM) in the indicated conditions. A Student’s t-test was used to compute statistical significance. Asterisks indicate statistical significance, whereas ns represents non-significance.

To gain deeper insight into the mode of action of these metabolites in inhibiting Ste2p signaling, we first analyzed their impact at the orthosteric site, i.e., α-factor binding. We performed protein-peptide docking of Ste2p and α-factor and observed that metabolite-binding at the Ste2p-Gpa1p interface favors the α-factor interaction, as inferred from the binding free energies ΔG (kcal/mol) **(Supplementary Figure 9b-c)**. We also analyzed protein-protein interaction between the Ste2p (GPCR) and miniGpa1p (55) by selecting GPCR configurations with and without metabolite-induced altered cavity topologies (IC4 and IC5), respectively (frames output from the aforementioned Ste2p-metabolite complex simulations), and computed the dissociation constant (Kd), binding affinity (ΔG), and the structural changes in the overall Ste2p topology **(Figure 3d-e, Supplementary Figure 9b-d)**. These computational analyses revealed that, in contrast to the metabolite-free Ste2p-Gpa1p interaction, referred to as the wild-type (WT) condition, the Kd value is many-fold lower in the presence of metabolites, indicating a cohesive response induced by these metabolites. A multi-fold lower Kd value further indicates and potentially explains that the metabolite binding favors the Ste2p (GPCR) and miniGpa1-protein interaction and enables the establishment of a stable complex that might influence the shielding of the effector-regulating domains of the Gpa1p or influence its binding with the Ste4p (Gβ)-Ste18p (Gγ) complex.

We next asked whether the observed resistance toward α-factor-induced PCD in single metabolic mutants might be the direct consequence of the identified metabolites or a pleiotropic response due to an altered genome. To test this, we exogenously supplemented wild-type yeast with zymosterol (ZST), ubiquinone 6 (CoQ6), and lanosterol (LST). Yeast cells were pre-loaded with metabolites for 24 hours at concentrations (0.1 μM, 1 μM, and 10 μM) that did not significantly alter growth profiles, except for 10 μM lanosterol, which showed mild perturbation (**Supplementary Figure 10a**), thereby ensuring the physiological relevance of our experimental conditions. Following pre-treatment, cells exhibited marked rescue from α-factor-induced PCD across multiple assays, including growth kinetics, PI-based viability, and FUN^TM^1 measurements **(Figure 3f-g, Supplementary Figure 10b-c)**. This protective effect was specific to α-factor-induced PCD, as metabolite-treated cells remained sensitive to acetic acid-induced death, suggesting that it is likely due to modulation of the Ste2p-Gpa1p interface and not a consequence of an altered response (**Figure 3g**). Next, we probed whether the observed modulation of Ste2 signaling also applies to the natural yeast mating behavior. We investigated this by performing a mating assay, which also revealed a decline in Ste2 signaling in metabolite-preloaded cells, suggesting the interlink between these metabolites and Ste2p signaling. Notably, in the case of CoQ6, we failed to observe any significant decline in the mating response, consistent with its instability observed in molecular dynamics simulations **(Figure 3h, Supplementary Figure 10d).**

Further, we evaluated the deactivation of the pathway at the MAPK signaling level by monitoring the Fus3 phosphorylation, where the α-factor-induced p-Fus3 levels were significantly suppressed by ZST and LST **(Figure 3i)** but not CoQ6 (data not shown). For final validation, we employed a P*_FUS1_*-eGFP-CYC1 transcriptional reporter system. Metabolite pretreatment significantly reduced eGFP-positive cells following α-factor stimulation compared to controls (**Figure 3j**), further demonstrating that the tested metabolites can inhibit α-factor-induced Ste2p signaling. These findings suggest endogenous metabolites as modulators of Ste2p signaling by stabilizing Ste2p-Gpa1p interactions.

### Site-Directed *Ste2* Mutants Abrogate Metabolite-Mediated Rescue Phenotype

To gain a deeper understanding, we investigated the role of metabolite binding in modulating Ste2p-Gpa1p interaction dynamics and employed both computational and experimental approaches. First, we conducted *in silico* screening by generating site-directed mutants of Ste2p designed to alter metabolite binding, followed by docking with Gpa1p to assess the impact of these mutations. We selected mutation sites by analyzing non-covalent interactions in Ste2p-metabolite complexes (IC4 and IC5). We prioritized stronger hydrogen bonds over weaker hydrophobic interactions and pinpointed specific interaction sites for CoQ6 (S75, R233), ZST (L289), and LST (T155, V152, I153). We finally applied a 2.7 Å to 3.3 Å ideal distance range for hydrogen bonds and selected mutants S75A for CoQ6, L289K for ZST, and T155D for LST. This filtering step eliminated unstable or distorted hydrogen bonds. These mutations significantly increased the dissociation constant (Kd) of the Ste2p-Gpa1p complex, indicating weakened interactions compared to wild-type Ste2p **(Figure 4a-b, Supplementary Figure 11a)**. This computational evidence supports the hypothesis that metabolite binding stabilizes the Ste2p-Gpa1p complex, facilitating the rescue response to α-factor-induced programmed cell death (PCD).

**Figure 4:**
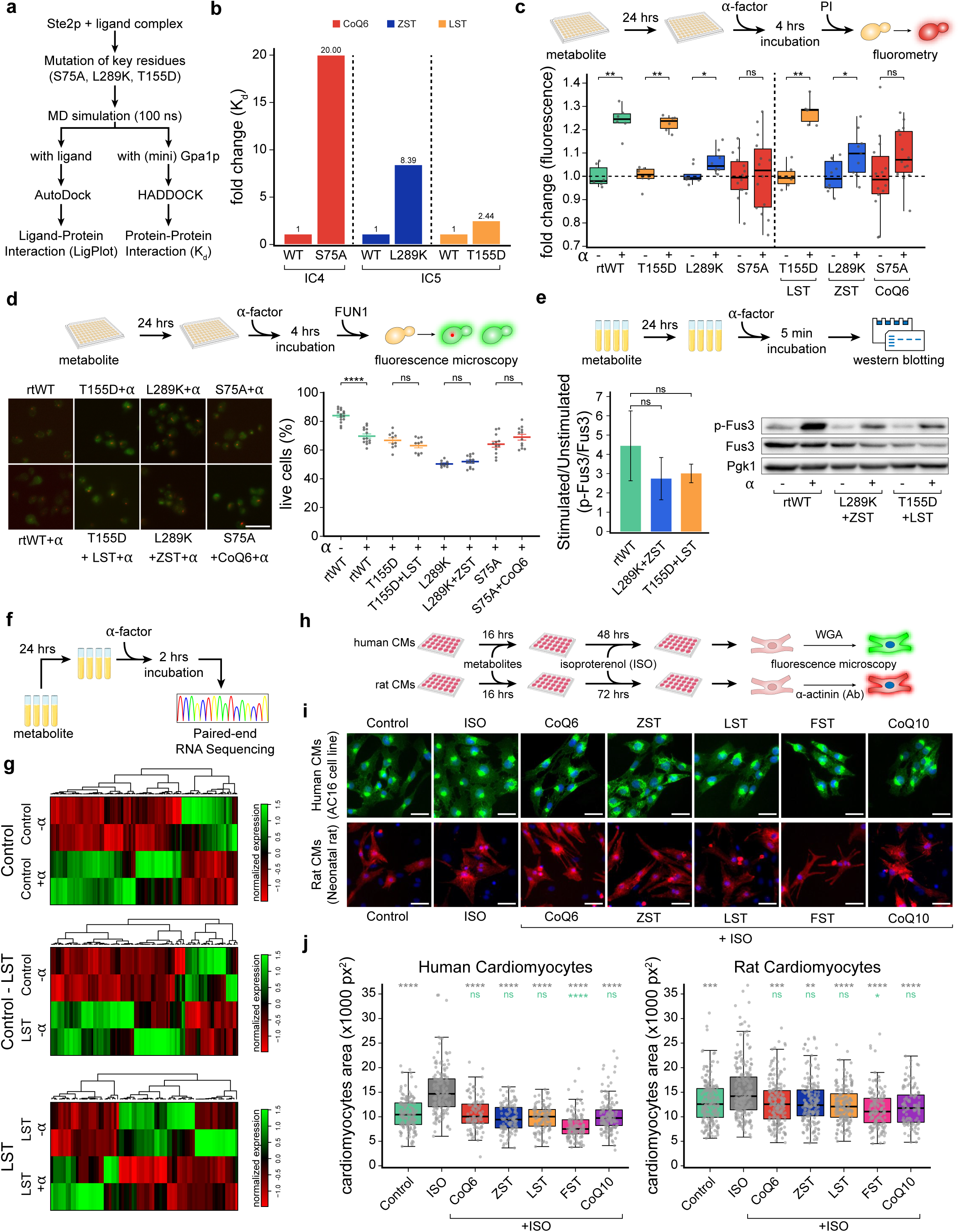
Site-Directed Ste2p Mutants Disrupt Metabolite-Mediated Rescue. **(a)** Workflow depicting the steps involved in Ste2p-miniG-protein docking of the wild-type and site-directed Ste2p mutants. Notably, docking was performed using HADDOCK and PRODIGY web servers. **(b)** Barplots depicting the dissociation constant (K_d_) fold change in Ste2p site-directed mutants and wild-type. Notably, fold change was computed with respect to the metabolite-influenced wild-type condition (Ste2p-miniG-protein). **(c)** The schematic diagram depicts the experimental workflow used to quantify α-factor-induced programmed cell death in generated site-directed missense mutants (T155D, L289K, S75A), alongside reconstituted wild-type *STE2* (rtWT), using a propidium iodide-based cell viability fluorometric assay. The box plot (left) depicts the increase in the relative proportion of dead cells upon α-factor exposure. Box plot (right) depicting the loss of rescue phenotype from the α-factor-induced programmed cell death in the indicated conditions when pre-loaded with metabolites as inferred using propidium iodide-based cell viability fluorometric assay. The y-axis represents the fold change of the propidium iodide fluorescence values with respect to their respective controls. The Mann-Whitney U test was used to calculate statistical significance. Asterisks indicate statistical significance, whereas ns represents non-significance. **(d)** Schematic representation (top) of the experimental approach used to measure cell vitality and viability using microscopy-based FUN^TM^1 staining. Representative micrographs (below) depicting the FUN^TM^1 staining results in the indicated conditions, Scale 10 µm. Mean-whisker plot depicting the relative proportion of the vital and viable yeast cells observed using FUN^TM^1 staining in the indicated conditions (n = 4 biological replicates). A Student’s t-test was used to compute statistical significance. Asterisks indicate statistical significance, whereas ns represents non-significance. Error bars represent the standard error of the mean (SEM). **(e)** Schematic representation (up) depicting the experimental design of phospho-MAPK activity-based Western blot. Barplots (down) depicting the p-Fus3 levels (mean ± SEM) in the indicated conditions. The y-axis represents the p-Fus3/Fus3 ratio for the stimulated condition normalized by its corresponding unstimulated sample. A Student’s t-test was used to compute statistical significance. Asterisks indicate statistical significance, whereas ns represents non-significance. **(f)** Schematic representation depicting the experimental design of RNA sequencing, featuring treatment duration and the sequencing parameters. **(g)** Heatmap depicting the expression of differentially expressed genes obtained from RNA sequencing in the indicated conditions. Notably, Control and LST represent yeast cells unloaded and pre-loaded with lanosterol, respectively. α-factor is represented as α, where plus and minus signs represent its presence and absence, respectively. **(h)** Schematic representation of the experimental workflow followed to deduce the impact of indicated metabolites treatment on isoproterenol (ISO)-induced, GPCR-mediated hypertrophy response in human (AC16) and neonatal rat cardiomyocytes. Notably, in the case of AC16 cells, Wheat germ agglutinin (WGA) was used to stain the cardiomyocytes, whereas, for neonatal cardiomyocytes, alpha-sarcomeric actinin staining was used. **(i)** Micrographs depicting the human (above; green colored) and neonatal rat (below; red colored) cardiomyocytes in the indicated conditions. Scale 50 µm. **(j)** Box plots depicting the surface area of human (AC16) and neonatal rat cardiomyocytes in the indicated conditions. Statistical significance of indicated metabolites with untreated control and isoproterenol-treated conditions is indicated in green and grey text, respectively. Mann-Whitney U test with Bonferroni-corrected p-values was used to compute statistical significance.

Next, we performed experimental validation by generating site-directed missense mutants targeting key binding residues at the Ste2p-Gpa1p interface and confirmed these computational predictions **(Supplementary Figure 11b, Supplementary Information)**. These mutants were expressed in a *ste2Δ* background, with the reconstituted wild-type *STE2* (rtWT) serving as a control. Using fluorometry-based cell death assays, we first assessed whether mutants retained the metabolite-mediated rescue response observed in wild-type cells. While rtWT exhibited significant cell death upon α-factor exposure that could be rescued by metabolite pretreatment, the T155D and L289K mutants showed no rescue response despite pretreatment with their respective metabolites (**Figure 4c**). This loss of rescue suggests direct metabolite regulation of Ste2p signaling at the intracellular Ste2p-Gpa1p interface. Notably, S75A mutants showed minimal α-factor responsiveness overall, likely due to significant structural disruptions affecting both pheromone sensitivity and metabolite binding. FUN^TM^1 staining assay and p-Fus3 signaling analysis by mapping MAPK pathway activation further supported these findings (**Figure 4d-e**). The lack of rescue highlights the direct role of metabolite binding at the Ste2p-Gpa1p interface in regulating downstream signaling. Interestingly, S75A mutants showed no α-factor-induced effects, likely due to significant structural disruptions. Shmoo formation assays further corroborated these findings, with no rescue effects observed in the mutants despite metabolite pre-loading **(Supplementary Figure 11c-e)**.

To gain an unbiased view of the mode of action of these metabolites in attenuating Ste2p (GPCR)-mediated pheromone-induced cell death in yeast, we performed RNA sequencing on LST-preloaded and untreated (control) cells with and without α-factor exposure **(Figure 4f)**. Transcriptomic analysis of control cells (without metabolite treatment) revealed significant expression changes in genes related to critical cellular processes upon α-factor exposure **(Figure 4g, top panel; Supplementary Figure 12c)**. A more detailed and careful examination revealed differential expression of genes implicated in PCD and mating responses, including *GSY1*, whose downregulation was linked to α-factor resistance **(Figure 4f-g, Supplementary Figure 12a-c, Supplementary Tables 7-10)**. LST treatment alone (without α-factor) also induced significant differential gene expression associated with cellular processing, validating its bioactivity **(Figure 4g, middle panel)**. However, comparison of LST-preloaded versus control cells following α-factor treatment revealed no prominent mating or PCD-related genes among the differentially expressed genes **(Figure 4g, lower panel)**; however, one cannot rule out the contribution of these genes in providing innate resistance to the α-factor. We tested the contribution of these genes in facilitating metabolite-mediated rescue phenotypes using the α-factor-induced cell-death assay on the knockouts of these differentially upregulated transcripts. Our results showed a significant loss of metabolite-mediated rescue phenotype in six out of ten knockouts, with *YCR095W-A* displaying the most pronounced phenotype loss **(Supplementary Figure 12d-f)**. These results collectively suggest that metabolite binding at the Ste2p-Gpa1p interface directly drives rescue responses, with secondary contributions from differentially expressed genes in attenuating α-factor-induced cell death.

Since our intensive computational interrogation of all the available human GPCR-Gα complexes revealed a higher degree of functional conservation **(Figure 1k)**, we next explored whether intracellular allosteric modulators such as ubiquinone 6 (CoQ6), zymosterol (ZST), lanosterol (LST), fucosterol (FST), and ubiquinone 10 (CoQ10) modulate GPCR signaling in higher vertebrates such as human and rat beta 1/2-adrenergic receptors signaling. Briefly, by using Gcoupler, we identified the putative GPCR-Gα interface and performed molecular docking with the aforementioned metabolites. Docking results revealed a high binding affinity of these selected metabolites at the GPCR-Gα interface of adrenergic receptors, reminiscent of Ste2p-metabolite interactions **(Supplementary Figure 13a-b)**. Sequence conservation analysis of the GPCR-Gα interface across yeast Ste2p and adrenergic receptors in humans and rats further confirmed a high degree of evolutionary conservation at the metabolite-binding residues **(Supplementary Figure 13c)**. Finally, to test this functional relevance, we evaluated the effect of these metabolites on isoproterenol-induced adrenergic receptor-mediated cardiac hypertrophy in human AC16 cardiomyocytes and neonatal rat cardiomyocytes. Preloading cells with these metabolites significantly attenuated hypertrophic responses, as evidenced by reduced single-cell surface area in quantitative assessments **(Figure 4h-j)**. Notably, to further evaluate the evolutionary conservation of this phenomenon, we also analyzed 75 unique GPCR-Gα complex structures from six species, selected from the PDB database. Dynamic docking was performed using five metabolites (CoQ6, ZST, LST, FST, and CoQ10) identified by Gcoupler as potential allosteric modulators and five negative controls predicted as poor binders. Results revealed significantly lower docking scores (<-7 kcal/mol) for Gcoupler-recommended metabolites compared to negative controls, irrespective of GPCR type or species **(Supplementary Figure 13d-h)**. These findings demonstrate that intracellular metabolite modulation of GPCR activity is a conserved mechanism extending beyond yeast to higher vertebrates.

## DISCUSSION

Over the last few decades, extensive research has focused on identifying allosteric modulators of GPCRs due to their relevance in drug discovery (Leach et al., 2007; Topiol, 2018a). Most known modulators are exogenous and target extracellular sites, while intracellular allosteric sites, identified recently through structural biology, offer novel avenues for regulation (Calebiro et al., 2021; Leach et al., 2007; Rees et al., 2002; Topiol, 2018a; van der Westhuizen et al., 2015; Wold and Zhou, 2018). These sites, overlapping with G-protein and β-arrestin coupling regions, highlight the potential for intracellular allosteric modulation (Bourque et al., 2022; van der Westhuizen et al., 2015; Yang et al., 2021). Intracellular modulators, including chemically diverse agents like auto-antibodies and sodium ions, remain poorly understood, emphasizing the need for systematic exploration of these sites (Bourque et al., 2022; Nachtergaele et al., 2012; O’Callaghan et al., 2012; Rees et al., 2002; Reyes-Alcaraz et al., 2020; Stornaiuolo et al., 2015; van der Westhuizen et al., 2015; Wold and Zhou, 2018; Zhang et al., 2015). However, the lack of data on intracellular modulators limits the feasibility of conventional computational approaches (Basith et al., 2018; Congreve et al., 2020; Hedderich et al., 2022)(Bartuzi et al., 2018; Chatzigoulas and Cournia, 2021; Hou et al., 2021; Topiol, 2018b).

To address this gap, we developed Gcoupler, a computational framework integrating *de novo* cavity identification, ligand synthesis, statistical analysis, graph neural networks, and Bioactivity-based ligand prioritization. Unlike existing tools, Gcoupler does not require cavity-specific experimentally validated compounds for model training. Gcoupler’s precision in cavity mapping, flexibility for user-defined queries, and ability to screen large chemical libraries make it a versatile and efficient tool. Additionally, Gcoupler’s generic design allows application beyond GPCRs, contrasting with existing platforms that often have limitations in modularity, precision, or open-source availability. Noteworthy, in contrast to other known allosteric sites identification tools for GPCRs, such as Allosite (Huang et al., 2013), AllositePro (Song et al., 2017), AlloReverse (Zha et al., 2023), that largely leverage the ML-based models or require orthosteric ligand-bound structure as input, the cavity detection feature of Gcoupler (LigBuilderV3), a critical step for the entire workflow, is not limited to only the allosteric sites; instead, it identifies all possible cavity-like regions on the protein surface, which then gets classified into druggable, undruggable, or amphibious based on their individual scoring and ligandability, thus making it unbiased and more specific towards query protein. Notably, the rationale for opting for LigBuilder V3 for cavity identification over similar tools such as Fpocket (Le Guilloux et al., 2009) is that the former uses a hydrogen atom probe, moving along the protein surface grid of 0.5 Å for cavity detection, being much more precise in detecting cavity boundaries, in both breadth and depth mapping; in contrast, the latter considers clusters of alpha spheres **(Supplementary Table 11)**.

To date, only a few methods leverage generative AI models for cavity/pocket-based drug design. Gcoupler is an open-source, end-to-end platform integrating Ligand-Based Drug Design (LBDD) and Structure-Based Drug Design (SBDD) for drug design and large-scale screening. Unlike Pocket Crafter (Shen et al., 2024), which requires proprietary tools (e.g., MOE QuickPrep) and lacks predictive model-building modules, Gcoupler offers comprehensive functionality. Similarly, DeepLigBuilder (Li et al., 2021) and Schrodinger’s AutoDesigner are either closed source or limited in features compared to Gcoupler. Comparative analysis highlights Gcoupler’s unique advantages in precision, flexibility, and functionality **(Supplementary Table 11)**.

Using Gcoupler, we investigated the molecular basis of innate resistance to α-factor-induced programmed cell death (PCD) in yeast. Unlike humans, yeast possess only two GPCR systems, making their pheromone-sensing pathway ideal for focused study. Previous research predominantly identified downstream regulatory mechanisms (Alvaro and Thorner, 2016; Sokolov et al., 2020; Velazhahan et al., 2021); however, our findings suggest an upstream, receptor-level regulation via endogenous intracellular metabolites. Computational and experimental evidence pinpointed specific metabolites binding to the Ste2p-miniGpa1 interface, modulating signaling. Site-directed mutagenesis confirmed the functional relevance of these metabolite-interacting residues. Of note, previous mutagenesis experiments also revealed multiple critical amino acid residues that overlap with IC4 and IC5, suggesting their functional relevance in Ste2p downstream signaling **(Supplementary Table 12)**. Despite these advances, certain limitations remain, including the potential pleiotropic effects in metabolic gene knockouts and challenges in replicating natural metabolite concentrations. Mechanistic insights into sterol biosynthesis mutants (*ergΔ*) revealed impaired mating responses due to heterogeneous defects, such as reduced sterol accumulation and shmoo formation, impaired membrane fusion, and decreased *FUS1* expression (Aguilar et al., 2010; Bagnat and Simons, 2002; Heese-Peck et al., 2002; Jin et al., 2008; Tiedje et al., 2007). This highlights a novel role for sterols in GPCR regulation and their broader implications for yeast microbial factories and stress tolerance (Ostrov et al., 2017; Shaw et al., 2019). Sterols and other endogenous metabolites were shown to modulate GPCR activity by targeting conserved Gα-binding sites, reinforcing the evolutionary conservation of this mechanism across species, as demonstrated in human and rat hypertrophy models in this study.

In summary, our work uncovers a novel regulatory mechanism for GPCRs mediated by intracellular metabolites and presents a computational framework, Gcoupler, to explore unexplored allosteric sites. The proposed model suggests that selective metabolites binding to GPCR-Gα interfaces induce local conformational changes, stabilizing GPCR-G-protein complexes and potentially obstructing downstream signaling. Alternative mechanisms, such as orthosteric site modulation, kinase/arrestin interaction interference, or alterations in membrane dynamics, remain to be explored, warranting further investigation. A critical limitation of our study is the absence of direct binding assays to validate the interaction between the metabolites and Ste2p. While our results from genetic interventions, molecular dynamics simulations, and docking studies strongly suggest that the metabolites interact with the Ste2p-Gpa1 interface, these findings remain indirect. Direct binding confirmation through techniques such as surface plasmon resonance, isothermal titration calorimetry, or co-crystallization would provide definitive evidence of this interaction. Addressing this limitation in future work would significantly strengthen our conclusions and provide deeper insights into the precise molecular mechanisms underlying the observed phenotypic effects. Another critical limitation of our findings is its reliance on tools like AutoDock and PRODIGY for preliminary binding affinity estimates, which lack the thermodynamic precision of advanced methods. Although calibrating docking scores with experimental data or using alchemical free energy calculations can improve accuracy, these methods are computationally expensive and require high-quality data, which is often unavailable. Gcoupler prioritizes speed, scalability, and accessibility, especially for data-sparse scenarios. By focusing on efficient, data-driven classification methods, it balances performance with practicality for large-scale screening. In this study, to address this limitation, we employed MD simulations with MM/GBSA, incorporating factors like protein flexibility and solvation effects for more accurate ΔG calculations. While computationally intensive approaches were beyond this study’s scope, we ensured reported ΔG values reflected system conformational flexibility by basing them on pre-simulated docked structures from molecular dynamics simulations. Further, our results suggest that the metabolite binds to the Ste2p-Gpa1 interface and modulates receptor activity upon pheromone stimulation, as supported by various assays. However, the precise sequence of interactions between Ste2p, the metabolite, and Gpa1 remains unexplored, as it requires sequential experiments beyond this study’s scope. Taken together, addressing these limitations in future work would significantly strengthen our conclusions and provide deeper insights into the precise molecular mechanisms underlying the observed phenotypic effects.

## MATERIAL AND METHODS

### Backend code for the Gcoupler

The back-end code for Gcoupler is implemented entirely in Python (3.8) and comprises four modules: Synthesizer, Authenticator, Generator, and BioRanker. Synthesizer employs LigBuilder V3.0 (Yuan et al., 2020) for *de novo in silico* ligand synthesis, identifying protein cavities likely to be active or allosteric sites using a hybrid GROW-LINK approach with a Genetic Algorithm. The module autonomously selects one cavity for ligand synthesis based on user-defined residue positions. Using the CAVITY function of LigBuilder (Yuan et al., 2011), it classifies 3D grid points around the protein into occupied, vacant, and surface points and integrates geometric and physicochemical properties to identify binding sites (Clark et al., 1989). Synthesizer outputs ligand structures in SMILES and PDBQT formats, alongside cavity grid coordinates for downstream modules. Authenticator validates the synthesized ligands using AutoDock Vina (1.2.3) for virtual screening (Trott and Olson, 2010). Binding energy calculations classify ligands into High-Affinity Binders (HABs) and Low-Affinity Binders (LABs), preserving balance for subsequent deep learning analysis. The default energy threshold is set to −7 kcal/mol, but users can explore alternative cutoffs, visualize distributions, and perform statistical comparisons within the workflow. These steps ensure precise identification and prioritization of ligand candidates for further analysis. The Authenticator uses the Kolmogorov-Smirnov test (Berger and Zhou, 2014), Anderson-Darling test (Engmann and Cousineau, 2011), and Epps-Singleton test (Goerg and Kaiser, 2009) for hypothesis testing for the comparison of the distributions. The Authenticator module visualizes ligand distributions using overlapping density plots and Empirical Cumulative Distribution Function (ECDF) curves. If the default threshold fails to produce statistically meaningful separation, users can supply alternative negative datasets, such as decoys generated via Gcoupler’s inbuilt RDKit Chem module or custom datasets (Landrum, n.d.). The Generator module builds deep learning-based classification models using the DeepChem (2.6.1) library (Ramsundar et al., 2019). It accepts High-Affinity Binders (HABs) and Low-Affinity Binders (LABs)/decoys from the Authenticator module to train four graph-based models: Graph Convolution Model (GCM), Graph Convolution Network (GCN) (Kipf and Welling, 2016), Attentive FP (AFP) (Xiong et al., 2020), and Graph Attention Network (GAT) (Veličković et al., 2018). Class imbalance is addressed through upsampling techniques. Generator tests all models using default hyperparameters, returning performance metrics for user selection. Hyperparameters can be tuned via manual settings or using default values followed by k-fold cross-validation. The final optimized model, trained on the complete synthetic dataset (HAB + LAB/decoys), enables large-scale screening of user-supplied compounds based on their SMILES representations. The BioRanker module performs post-prediction analysis for functional activity-based compound screening. Positively predicted compounds are selected using a stringent probability threshold or adaptive methods such as G-means and Youden’s J statistic, which optimize sensitivity and specificity. The selected compounds are projected into biological activity spaces (Chemistry, Targets, Networks, Cells, Clinics) by comparing their biological activity descriptor vectors with those of HABs using cosine similarity (Bertoni et al., 2021). A modified PageRank algorithm ranks compounds based on activity-specific scores, with support for multi-activity ranking to refine results based on user-defined biological properties, ensuring precise and context-relevant compound prioritization.

Additional information about the backend code for Gcoupler, along with methodology for Runtime analysis, Sequence-structural-functional level analysis, Molecular Dynamics Simulation, Molecular docking (AutoDock), Functional enrichment analysis, and Protein-protein docking, can be accessed in the **Supplementary Information**.

### Gcoupler Benchmarking

To assess batch effects across Gcoupler runs for a specific cavity, we utilized the standard Gcoupler Docker image. Intracellular cavity 4 (IC4) of the Ste2 protein of yeast was used for benchmarking. A total of 100 molecules were *in silico* synthesized by the Synthesizer module of Gcoupler iteratively. Post-generation, atom pair fingerprints (ChemmineR; R package) were calculated for the synthesized molecules from each run, and the data was visualized using Principal Component Analysis and pairwise comparison using Tanimoto Similarity (ChemmineR, R package).

For model benchmarking, Gcoupler was validated on GPCRs from the DUD-E dataset, alongside the information about the active ligands and their randomly selected number-matched decoys (Mysinger et al., 2012). Additionally, Gcoupler’s performance in identifying experimentally elucidated allosteric sites and modulators was tested using PDB complexes obtained from the RCSB PDB database.

### Metabolomics

Wildtype (BY4741) and *ste2Δ* yeast strains were grown in YPD medium at 30°C, 150 rpm, through primary and secondary cultures (16 hours each). Equal cell numbers (1.5 mL) were aliquoted into a 96-well deep well plate. α-factor (Sigma-Aldrich) was added at final concentrations of 10, 20, 30, 40, and 50 μM (eight replicates each). DMSO served as the solvent control, while untreated WT and *ste2Δ* conditions received no treatment. Plates were incubated (30°C, 150 rpm, 4 hours) under a breathable membrane. A 50 μL aliquot was taken for Propidium Iodide (PI; 11195, SRL) assay as described in **Supplementary Information**. Following the PI assay, four pooled replicates were pelleted (6000 rpm, 5 min, RT), treated with zymolyase (40 U/mL, 1X PBS, 30°C, 1 hour), washed with PBS, and metabolomics analysis was performed. Data analysis included peak normalization, omission of metabolites with constant or >50% missing values, and kNN-based imputation (MetaboAnalyst). Data were IQR-filtered, and differentially enriched metabolites (DEMs) were identified by calculating log2 fold change (|log2FC| ≥ 1, p < 0.05 via Student’s t-test). Pathway Over-Representation Analysis (ORA) was performed using MetaboAnalyst with hypergeometric or Fisher’s exact tests to assess pathway enrichment against background metabolite distributions. Further details about the methodology are available in **Supplementary Information**.

### Genetic Screening

Fifty-three knockout strains from the Yeast Deletion Collection, along with WT and *ste2Δ*, were treated with α-factor (30 μM), while DMSO served as the solvent control. Plates were incubated for 4 hours. A 50 μL aliquot was used to measure the Propidium iodide-based cell viability assessment assay as described in the Supplementary Information. Fluorescence data were normalized to blank-adjusted OD600, followed by two additional rounds of normalization with unstained and HK controls. The percentage fold change for the treated group was calculated relative to the untreated group, and statistical significance was determined using a one-sample Student’s t-test. Further details about the methodology used is available in **Supplementary Information**.

### Pre-loading of Yeast cells with a metabolite

Yeast cells were cultured in YPD medium at 30°C, 200 rpm for 16 hours in primary and secondary cultures. Equal cell densities (5 μL) from secondary cultures were inoculated into 96-well plates containing 145 μL YPD with metabolites Coenzyme Q6 (CoQ6, 900150O, Avanti^®^ Polar Lipids), Zymosterol (ZST, 700068P, Avanti^®^ Polar Lipids), and Lanosterol (LST, L5768, Sigma-Aldrich) at 0.1 μM, 1 μM, and 10 μM concentrations. Plates were incubated for 24 hours at 30°C, 200 rpm, with multiple biological replicates. Ethanol-treated wells served as solvent controls. For site-directed *STE2* mutants, the mutants were grown in YPD for primary and secondary cultures, but the metabolite pre-loading was performed in YPGR instead of YPD to induce Ste2 expression.

After pre-loading, the following assays were performed: growth kinetics, propidium iodide-based assay, FUN^TM^1 staining, Mating assay, phospho-MAPK activity-based western blot, and transgenic reporter assay. The detailed protocol for each of these assays is available in **Supplementary Information**.

### RNA-Sequencing

Yeast cells (BY4741) were cultured in YPD medium at 30 °C, 200 rpm, with or without lanosterol (LST, 1 μM) in biological duplicates. Cells were subsequently treated or untreated with α-factor (30 μM) for 2 hours. RNA was isolated following Mittal et al. (Mittal et al., 2022). Sequencing quality was assessed using MultiFastQ, and paired-end reads were trimmed and aligned to the S. cerevisiae reference genome (ENSEMBL R64-1-1; GCA_000146045.2) using the Rsubread package (v2.6.4). Gene expression counts were generated via featureCounts and normalized using TMM **(Supplementary Table 7)**. Differential expression analysis was performed with NOISeq (v2.38.0) **(Supplementary Tables 8–10)**. Functional enrichment was assessed using the Gene Ontology Term Finder (v0.86) from the Saccharomyces Genome Database. Raw FASTQ files and normalized expression data are available on Zenodo. Additional details are available in the **Supplementary Information**.

### Site-Directed Mutagenesis

The gene encoding wild-type *STE2* was PCR amplified from the genome of *Saccharomyces cerevisiae* and further cloned into plasmid pRS304 under galactose-inducible *GAL1* promoter to generate plasmid pRS304-P*_GAL1_*-STE2-CYC1 using Gibson assembly (Gibson et al., 2010, 2009). The mutants were generated by PCR amplifying the gene with primers consisting of respective mutations and cloned into plasmid pRS304 under *GAL1* promoter to generate pRS304-P*_GAL1_*-STE2 S75A-CYC1, pRS304-P*_GAL1_*-STE2 T155D-CYC1, and pRS304-P*_GAL1_*-STE2 L289K-CYC1. Wild-type (rtWT) and mutant *STE2* were integrated by digesting the pRS304 vector with restriction enzyme BstXI to generate a linearized plasmid and transformed into the *Saccharomyces cerevisiae* BY4741 *ste2Δ* strain. Additional details about the methodology are available in the **Supplementary Information**

### Cardiomyocytes Hypertrophy Models

Human AC16 cardiomyocytes were cultured in DMEM-F12 (Thermo Scientific) with 12.5% fetal bovine serum (FBS) at 37°C and 5% CO2. Cells were seeded in a 24-well plate for size measurements, treated after 24 hours with metabolites (CoQ6, ZST, LST, FST, CoQ10) at 2.5 μM, and incubated overnight with 1% FBS. The medium was refreshed with fresh metabolites and isoproterenol (25 μM) for 48 hours. Cells were washed with PBS, fixed with 4% paraformaldehyde, and stained with wheat germ agglutinin (Thermo Scientific) and DAPI. Images were captured using a Leica DMI 6000 B microscope at 20X magnification, and cell area was measured using ImageJ. Neonatal rat cardiomyocytes were isolated from 1-3-day-old SD rat pups using Collagenase Type II. After heart explantation and digestion, the cells were centrifuged and pre-plated for 90 minutes to remove fibroblasts. The cardiomyocytes were seeded in a gelatin-coated 24-well plate, incubated overnight with 2.5 μM metabolites and 1% FBS, and then treated with metabolites (2.5 μM) and isoproterenol (10 μM) for 72 hours. Cells were fixed, stained with alpha-sarcomeric actinin and DAPI, and images were captured using a Leica DMI 6000 B at 20X magnification. Cell area was quantified using ImageJ. Additional details about the methodology are available in **Supplementary Information**.

### Statistical Analysis

Statistical analyses were performed using Past4 software or R Programming. The Mann-Whitney U test was applied to compare medians between two distributions (non-parametric), while Student’s t-test was used for pairwise comparisons of means. P-value correction was performed using the Bonferroni method when necessary. A significance threshold of 0.05 was set, with *, **, ***, and **** indicating p-values <0.05, <0.01, <0.001, and <0.0001, respectively.

## Data Availability

The processed untargeted metabolomics data is provided as Supplementary Information. The raw RNA sequencing files are available at ArrayExpress under accession *E-MTAB-12992*.

## Code Availability

A Python package for Gcoupler is provided via pip https://test.pypi.org/project/Gcoupler/. A docker container pre-compiled with Gcoupler and all of its dependencies can be found at https://hub.docker.com/r/sanjayk741/gcoupler. The source code of Gcoupler is available on the project GitHub page: https://github.com/the-ahuja-lab/Gcoupler and also at Zenodo with DOI: 10.5281/zenodo.7835335, whereas the raw sequencing files can be accessed using DOI: 10.5281/zenodo.7834294.

## Ethical statement

The local IAEC (Institutional Animal Ethics Committee) committee at CSIR-Central Drug Research Institute approved all the animal experiments (IAEC/2020/38) following the guidelines of the Committee for the Purpose of Control and Supervision of Experiments on Animals (CPCSEA), New Delhi, Government of India.

## Acknowledgments

The authors thank the IT-HelpDesk team of IIIT-Delhi for assisting with the computational resources. We thank all the members of the Ahuja lab for their intellectual contributions at various stages of this project. We thank Prof. G.P.S Raghava for providing critical comments on our manuscript. We thank Dr. Arjun Ray for providing intellectual support. We also thank Dr. Martin Graef and Dr. Kaushik Chakraborty for sharing yeast strains and the NIPER Guwahati central facility for helping us with high-resolution metabolomics. The Ahuja lab is supported by the Ramalingaswami Re-entry Fellowship (BT/HRD/35/02/2006), a re-entry scheme of the Department of Biotechnology, Ministry of Science & Technology, Government of India, Start-Up Research Grant (SRG/2020/000232) from the Science and Engineering Research Board, and a research grant from IHUB Anubhuti (Project Grant/23) and an intramural Start-up grant from Indraprastha Institute of Information Technology-Delhi. The INSPIRE faculty grant from the Department of Science & Technology, India, funds the Sengupta lab. Gupta Lab is funded by the Ramalingaswami Re-entry Fellowship (BT/RLF/re-entry/14/2019) from the Department of Biotechnology, Government of India.

## SUPPLEMENTARY FIGURE LEGENDS

**Supplementary Figure 1: Gcoupler Benchmarking using experimentally validated orthosteric ligands of GPCRs**

**(a)** AUC-ROC curves depicting the optimal probability cutoff recommended by G-means and Youden J index algorithms, respectively. Notably, the optimal threshold is indicated by the black dot on the ROC curve. **(b)** Molecular representation depicting the indicated experimentally validated orthosteric sites of the AA2AR, ADRB1, ADRB2, CXCR4, and DRD3 receptors, with the zoom-in inlet on the right highlighting the ligand and the cavity topologies. **(c)** Overlapping density plots comparing the distributions of synthetic compounds predicted to target the indicated receptors into High-Affinity Binders (HAB) and Low-Affinity Binders (LAB) using the Authenticator module of the Gcoupler package. **(d)** Bar graphs indicating the performance of the indicated models made for the experimentally elucidated orthosteric sites of the indicated GPCRs, generated using the Gcoupler workflow. **(e)** Confusion matrices indicating the relative proportions of experimentally determined ligands and their respective decoys (randomly selected to match # of ligands for each receptor). Note that the number of active ligands and decoys is mentioned at the bottom.

**Supplementary Figure 2: Gcoupler Benchmarking using experimentally validated allosteric ligands of GPCRs**

**(a)** Molecular representation of the β2AR, CCR2, and CCR9 receptors with highlighted experimentally validated intracellular allosteric cavity and ligand. **(b)** Overlapping density plots comparing the distributions of synthetic High-Affinity Binders (HAB) and Low-Affinity Binders (LAB) predicted to target the allosteric cavities of the indicated GPCRs using the Gcoupler package. **(c)** Bar graphs depicting the performance metrics of the indicated classification model generated using Gcoupler for the highlighted allosteric cavities of the indicated proteins.

**Supplementary Figure 3: Run Time comparison of Gcoupler and AutoDock**

**(a)** Flowchart depicting the entire workflow used to compute and compare the runtime using AutoDock and Gcoupler. Of note, the Synthesizer, Authenticator, and Generator module of Gcoupler are indicated alongside their key processes. **(b)** Ribbon diagram depicting the ADRA1A predicted structure using Alphafold (ID: AF-P35348). **(c)** A waterfall chart depicts the timestamp information about the key steps involved in AutoDock. Time consumed at the indicated steps is mentioned in seconds and hours along the y-axis and above bars, respectively. **(d)** Waterfall chart depicting the timestamp information about the key steps involved in Gcoupler. Time consumed at the indicated steps is mentioned in seconds and hours along the y-axis and above bars, respectively. **(e)** Bar plot depicting the comparison between the total time consumed by the AutoDock and Gcoupler to predict the binding of experimentally validated ligands for ADRA1A (AF-P35348). **(f)** Boxplot depicting the performance metrics of the 10-fold cross-validation of the training data obtained using Gcoupler. **(g)** Boxplot depicting the performance metrics of the 10-fold cross-validation of the testing data obtained using Gcoupler. **(h)** Scatterplot depicting the relationship between AutoDock computed binding energies and Gcoupler computed prediction probabilities of the experimentally validated ligands of ADRA1A protein. **(i)** Density plot depicting the distribution of RMSD between GPCR-Gα cavity across selected GPCRs-G-protein pairs. Of note, the RMSD of the Gα-protein cavity was normalized with the RMSDs of the respective whole proteins across all pairwise comparisons. **(j)** Heatmap depicting the RMSD values obtained by comparing all the GPCR-Gα-protein interfaces of the available human GPCRs from the protein data bank. **(k)** Heatmap depicting the RMSD values obtained by comparing the selected human GPCRs available in the protein data bank. **(l)** Density plot depicting the distribution of cosine similarities among the *in silico* synthesized ligands of the GPCR-Gα-protein interfaces of the available human GPCRs using Gcoupler.

**Supplementary Figure 4: Ste2 protein cavities prediction using Gcoupler**

**(a)** Schematic diagram depicting the hypothesis that the intracellular yeast metabolites could allosterically modulate the GPCR-Gα-protein (Ste2p-Gpa1p) interface and, therefore, the Ste2p signaling pathway in yeast. **(b)** Overlapping Ste2p (GPCR) ribbon diagram depicting the structural similarity between the experimentally elucidated Ste2p structure before and after molecular dynamics simulation using GROMACS software. RMSD value is depicted at the bottom. **(c)** Flowchart depicting the key steps used for the molecular dynamics simulation of Ste2p using CHARMM-GUI and GROMACS software. **(d)** Line plot depicting the changes in the density (kg/m³) along the z-axis of the three-dimensional Ste2p structure. **(e)** Line plot depicting the decline in the potential (kJ/mol) over molecular dynamics simulation time in picoseconds (ps). **(f)** Line plot depicting the changes in the density (kg/m³) of the overall system over molecular dynamics simulation time in picoseconds (ps). **(g)** Line plots depicting the changes in the pressure (in bars) and temperature (in Kelvin) of the overall system during molecular dynamics simulation of Ste2 protein. **(h)** Line plot depicting the Root Mean Square Deviation (RMSD) changes of the Ste2 protein during molecular dynamics simulation. Note that the RMSD values are indicated in nanometers (nm). **(i)** Ramachandran plot depicting the location of simulated Ste2 protein residues in the favorable and unfavorable regions. **(j)** Barplot depicting the predicted cavity volumes (Å). **(k)** Scatterplot depicting the relationship between the cavity volume and surface area in the indicated cavities of the Ste2 protein. **(l)** Heatmap depicting the percentage of amino acids of the indicated cavities residing in the EC (extracellular), IC (intracellular), and TM (transmembrane) region of the Ste2 protein. **(m)** Scatterplot depicting the predicted drug score of the indicated cavities of the Ste2 protein. **(n)** Snake plot depicting the key amino acids of the Ste2 protein alongside their location within the indicated functional cavities. **(o)** Snake plot depicting the conservation of Ste2 protein at the amino acid level. Of note, Ste2 proteins from 15 related yeast species were used for computing the conservation score.

**Supplementary Figure 5: Comparison of the chemical composition of cavity-specific ligands of Ste2 protein.**

**(a)** Venn diagram depicting the number of overlapping amino acids constituting the Extracellular Cavity 1 (EC1), Intracellular Cavity 4 (IC4), and Intracellular Cavity 5 (IC5) of the Ste2p. **(b)** Two-dimensional and **(c)** three-dimensional Principal Component Analysis plots depicting the segregation of *in silico* synthesized ligands for the indicated cavities via Gcoupler. Of note, atom pair fingerprints were used as features for this analysis. **(d)** Scatter plot depicting the relationship between the depth of the pharmacophore features and the neighboring grids. **(e)** Donut charts illustrating the properties of amino acids constituting the pharmacophore for each of the indicated cavities. **(f)** Two-dimensional and **(g)** three-dimensional Principal Component Analysis plots depicting the chemical properties of *in silico* synthesized ligands from five independent runs on the IC4 of the Ste2p using the Gcoupler Synthesizer module. Of note, atom pair fingerprints were used as features for this analysis. **(h)** Heatmap depicting the Tanimoto Similarities scores between the synthesized ligands for IC4 of Ste2p across five independent runs using Gcoupler.

**Supplementary Figure 6: Sanity Check of Gcoupler Predictions.**

**(a)** Principal Component Analysis depicting chemical heterogeneity of synthetic compounds synthesized using Gcoupler across intracellular cavities (IC4 and IC5). Notably, the compounds were segregated into High-Affinity Binders (HAB) and Low-Affinity Binders (LAB) by the Authenticator module of the Gcoupler. **(b)** Heatmap depicting the chemical similarity of the *de novo* synthesized HAB (dark green) and LAB (light green) across intracellular cavities (IC4 and IC5) of the Ste2 protein. Tanimoto similarities were computed using Atomic fingerprints. **(c)** Heatmaps illustrating the relative enrichment of the indicated functional groups (RNH2: primary amine, R2NH: secondary amine, R3N: tertiary amine, ROPO3: monophosphate, ROH: alcohol, RCHO: aldehyde, RCOR: ketone, RCOOH: carboxylic acid, RCOOR: ester, ROR: ether, RCCH: terminal alkyne, RCN: nitrile) among the Gcoupler-based *de novo* synthesized binders of Ste2p receptor across IC4 and IC5. **(d)** Box plots depicting the model’s performances across the 10-fold cross-validation. The upper and lower box plots depict the performance metrics of models trained using the *de novo* ligands for intracellular cavities (IC4 and IC5), respectively, of the Ste2 protein. **(e)** Venn diagrams depicting the number of yeast metabolites from YMDB predicted to bind to the Ste2p-Gpa1p interface, predicted using Gcoupler and AutoDock. Of note, IC4 and IC5 represent intracellular cavities 4 and 5 of the Ste2 protein, respectively. **(f)** Principal Component Analysis depicting chemical heterogeneity of endogenous yeast metabolites predicted to bind to the intracellular cavities (IC4 and IC5) of the Ste2 protein. Notably, the compounds were segregated into High-Affinity Metabolites (HAM) and Low-Affinity Metabolites (LAM) by the Generator module of the Gcoupler. **(g)** Heatmap illustrating the relative enrichment of the indicated functional groups among the endogenous intracellular allosteric modulators (metabolites) of Ste2 receptor across IC4 and IC5. **(h)** Percentage stacked barplot depicting the relative enrichment of the indicated functional groups of the High-Affinity Metabolites (HAM) and Low-Affinity Metabolites (LAM) across intracellular cavities (IC4 and IC5) of the Ste2 protein. **(i)** Heatmap depicting PageRank score of the selected metabolites for IC4 and **(j)** IC5 with respect to different biological properties. **(k)** Box plots depicting the performance parameters of the indicated models generated using Gcoupler against the IC4 cavity of Ste2p. Note that the training and testing datasets were generated randomly (5 iterations) from the *in silico* synthesized ligands from Gcoupler. **(l)** Scatter plots depicting the segregation of High/Low-Affinity Metabolites (HAM/LAM) (indicated in green and red) identified using the Gcoupler workflow with 100% training data. Of note, models trained on lesser training data size (25%, 50%, and 75% of HAB/LAB) severely failed to segregate High-Affinity Metabolites (HAM) and Low-Affinity Metabolites (LAM) (along the Y-axis). The X-axis represents the binding affinity calculated using IC4-specific docking using AutoDock. **(m)** Bar plots depicting the correlation values obtained between the Gcoupler prediction probabilities and AutoDock computed binding energies in the indicated training data size. **(n)** Box plots depicting the distributions for binding energies of the High-Affinity Metabolites and Low-Affinity Metabolites computed using cavity-specific docking and full docking via Autodock for IC4 and IC5 of the Ste2p.

**Supplementary Figure 7: Untargeted metabolomics and genetic screening suggest an interlink between metabolites and Ste2p-mediated Programmed Cell Death.**

**(a)** Screenshot of the central yeast metabolism from the Kyoto Encyclopedia of Genes and Genomes (KEGG) indicating the key metabolic pathways (highlighted in red) harboring metabolic intermediates predicted as high-affinity binders of GPCR-Gα-protein (Ste2p-Gpa1p) interface by Gcoupler. **(b)** Venn diagram depicting the percentage overlaps of the mating-related genes reported in the literature and those used in the screening. The *STE2* was the only common gene. **(c)** Heatmap depicting the growth curve profiles of the wild-type, single metabolic mutants, and *ste2Δ* under optimal growth conditions. **(d)** Schematic diagram depicting the experimental workflow of the untargeted metabolomics experiment. Notably, the experiment involves the usage of increasing doses of α-factor to induce PCD and to enrich the surviving cells selectively. **(e)** Boxplot depicting the increase in the propidium iodide fluorescence upon the increasing concentration of α-factor treatment. Heat-killed (HK), untreated, and vehicle (DMSO-treated) were used as controls, whereas the increasing concentration of α-factor surviving wild-type cells was used as test conditions. The Mann-Whitney U test was used to calculate statistical significance. Asterisks indicate statistical significance, whereas ns represents non-significance. **(f)** Heatmap depicting the relative (de)enrichment of differentially enriched metabolites in the indicated conditions. Of note, four biological replicates per condition were used in the untargeted metabolomics. **(g)** Principal Component Analysis depicting the segregation of indicated samples based on their metabolomics profiles along the PC1, PC2, and PC3. **(h)** Correlation plot depicting Pearson’s correlation of the metabolome of the indicated samples. **(i)** Heatmap depicting the relative enrichment of all the detected metabolites in the indicated condition. Heat-killed (HK), untreated, and vehicle (DMSO-treated) were used as controls, whereas the increasing concentration of α-factor surviving wild-type cells was used as test conditions. **(j)** Venn diagram indicating that 38 predicted intracellular allosteric modulators of Ste2p were identified in the untargeted metabolomics profiling. **(k)** Scatterplot depicting the Pathway Over Representation Analysis (ORA) results of the differentially enriched metabolites (treated vs. vehicle control) identified using untargeted metabolomics.

**Supplementary Figure 8: Docking and Molecular Dynamics simulations suggest the stability of the Ste2-metabolite complex.**

**(a)** Ligplots depicting the interacting amino acid residues and atoms of the Ste2 protein and indicated metabolites for the intracellular cavities (IC4 and IC5). **(b)** Table indicating the Van der Waals (ΔE_vdw_), electrostatic (ΔE_elec_), polar solvation (ΔE_psolv_), and non-polar solvation (ΔE_npsolv_) energies along with net binding free-energy (ΔG_bind_) in kcal/mol of the three independent replicates of the indicated metabolites across intracellular cavities IC4 and IC5 of the Ste2p. Note the values in the table are provided as mean ± standard errors. **(c)** Line plot depicting the variation in the RMSD of the indicated metabolites bound to the intracellular cavity 4 (IC4) at the GPCR-Gα interface of the Ste2p, obtained using longer MD simulation runs. Notably, RMSD is provided in Angstroms (Å), whereas the simulation time is in nanoseconds (ns).

**Supplementary Figure 9: Divergent characteristics of site-directed missense mutants of *STE2* gene.**

**(a)** Line plots depicting the total energy decomposition of the individual amino acids of the Ste2p (three independent replicates) computed using MD simulations in IC4 (left) and IC5 (right). **(b)** Molecular representation of the Ste2 protein structures (blue) and the miniGpa1-proteins (cyan), with the highlighted interface residues (red and green) in the indicated conditions. **(c)** Scatterplot depicting the net binding free-energy ΔG (kcal/mol) of the Ste2p (with and without indicated metabolites) with miniG-protein (y-axis) and α-factor (x-axis). **(d)** Barplot depicting RMSD values of the Ste2 protein with indicated metabolite across intracellular cavities IC4 and IC5.

**Supplementary Figure 10: Exogenous supplementation of selective metabolites rescues α-factor-mediated Programmed Cell Death.**

**(a)** Growth curve profiles of the wild-type (control), *ste2Δ,* and wild-type yeast pre-treated with the indicated metabolites. **(b)** Schematic diagram (left) depicting the workflow opted to monitor α-factor-induced programmed cell death (PCD) in the wild-type yeast cells pre-loaded for 24 hours with ubiquinone 6 (CoQ6), zymosterol (ZST), and lanosterol (LST) at three different concentrations. Beanplots on the right depict the changes in the α-factor-mediated PCD in the indicated conditions. Notably, cell viability was assessed by computing the area under the curve (AUC) of the growth kinetics. The Mann-Whitney U test was used to calculate statistical significance. Asterisks indicate statistical significance, whereas ns represents non-significance. **(c)** Beanplot depicting the fold change in the α-factor-mediated programmed cell death, assessed using the propidium iodide-mediated programmed cell death in the indicated conditions. The Mann-Whitney U test was used to calculate statistical significance. Asterisks indicate statistical significance, whereas ns represents non-significance. **(d)** Representative micrographs (left) depicting the growth of the cells (mating response) pretreated with indicated metabolites in the indicated single drop-out medium (SC-Lys and SC-Met). Quantifications of the mating response with respect to the SC-Met control are indicated as a bar graph (mean ± SEM) on the right (n = 3 biological replicates, each with three technical replicates). The Student’s t-test was used to compute the statistical significance. Asterisks indicate statistical significance, whereas ns represents non-significance.

**Supplementary Figure 11: Effects of Ste2 Mutants on Shmoo Formation in Response to α-Factor.**

**(a)** Ligplots depicting the interacting amino acid residues of the mutant Ste2 proteins (mutation site indicated) and atoms of indicated metabolites at the intracellular cavities (IC4 and IC5). **(b)** Growth curve profiles of the reconstituted wild-type (rtWT), and indicated missense site-directed mutants. **(c)** Schematic representation depicting the overall experimental workflow designed for the quantitative assessment of the shmoos formation upon α-factor treatment. **(d)** Micrographs depicting the phase contrast images of the rtWT cells, pretreated with the indicated metabolites, with and without α-factor treatment, Scale 10 µm. Mean-Whisker plot (right) indicating the relative proportions of shmoos forming cells in the indicated conditions. **(e)** Micrographs depicting the phase contrast images of the indicated site-directed missense mutants of the *STE2* gene, pretreated with the indicated metabolites, with and without α-factor treatment, Scale 10 µm. Mean-Whisker plot (right) indicating the relative proportions of shmoos forming cells in the indicated conditions. The Student’s t-test was used to compute the statistical significance of **d** and **e**. Asterisks indicate statistical significance, whereas ns represents non-significance. Error bars represent the standard error of the mean (SEM).

**Supplementary Figure 12: RNA-Sequencing unveils genes involved in attenuating α-factor-induced cell death response.**

**(a)** A table highlighting the key differentially expressed genes in the indicated conditions reported to be involved in regulating yeast mating response. Arrowheads and their direction represent the direction of differential expression. **(b)** A table highlighting the key differentially expressed genes in the indicated conditions reported to be involved in regulating yeast PCD-like signaling response. Arrowheads and their direction represent the direction of differential expression. **(c)** M-D plots (left) from the NOIseq analysis highlighting the differentially expressed genes (colored) in the indicated conditions. Bar plots (middle) depicting the prominent Gene Ontologies in the indicated conditions. Venn diagram (right) depicting the overlaps between the differentially expressed genes and the genes used in biochemical assays in this study. **(d)** Venn diagram depicting the overlap between the differentially expressed genes obtained from the RNA-Seq analysis and the yeast knockout library. **(e)** Growth curve profiles of the wild-type and indicated knockouts from the yeast knockout library. **(f)** Box plot depicting the relative proportion of dead cells upon α-factor exposure in the indicated knockouts from the yeast knockout library as inferred using propidium iodide-based cell viability fluorometric assay (n=12 biological replicates from 2 experiments). The y-axis represents the fold change of the propidium iodide fluorescence values with respect to their respective controls. The Mann-Whitney U test was used to calculate statistical significance. Asterisks indicate statistical significance, whereas ns represents non-significance.

**Supplementary Figure 13: Evolutionary Conservation of GPCR-Gα interface.**

**(a)** Molecular representations depicting the topology of human and rat ADRB1 and ADRB2 receptors. GPCR-Gα interface cavities are color-coded at the intracellular sites. Of note, in the case of human ADRB1, the nomenclature of the two cavities detected at the interface includes ‘Cv’ (cavity) succeeded by the numerical number. **(b)** Barplots depicting the binding energies obtained by the docking of Human ADRB1 (7BVQ), Human ADRB2 (5X7D), Rat ADRB1 (P18090), and Rat ADRB2 (P10608) with the indicated metabolites at the GPCR-Gα interfaces. **(c)** Table depicting the conservation of yeast Ste2p-metabolite (ZST, CoQ6, and LST) interacting residues in the adrenergic receptors from humans and rats. **(d)** The schematic workflow illustrates the steps involved in measuring and comparing the sequential and ligand-interaction conservation of GPCRs across six different species. **(e)** Donut charts illustrating the GPCR count across different species. **(f)** Phylogenetic tree representing 75 unique GPCRs from six species. **(g)** Empirical cumulative distribution function (ECDF) plot depicting binding energy distribution of Gcoupler identified potential allosteric modulators along with five negative controls. **(h)** Heatmap depicting the pairwise d-values/p-values from the Kolmogorov-Smirnov statistical test under specified conditions. The lower triangle of the heatmap represents d-values, while the upper triangle shows p-values for the specified pair. The test provides two key outputs: the D-value, which represents the maximum difference between the cumulative distribution functions (CDFs) of the two groups, and the p-value, which indicates the statistical significance of the observed difference. Symbols *, **, ***, and **** refer to p-values <0.05, <0.01, <0.001, and <0.0001, respectively.

## SUPPLEMENTARY TABLES

**Supplementary Table 1:** Table contains information about the structure of the human GPCR-Gα-Protein complexes downloaded from the Protein Data Bank (PDB). It also contains information about the PDB IDs, Protein name, species information, whether it is used in this study or not, UniProt IDs, and other information.

**Supplementary Table 2:** The table contains information about the normalized Root Mean Square Deviation (RMSD) of the pairwise comparison of the GPCR-Gα-Protein interface (cavities) for the indicated GPCRs.

**Supplementary Table 3:** The table contains information about the chemical similarities of the *de novo* synthesized synthetic compounds of the indicated GPCR-Gα-Protein interface (cavities). Of note, the chemical similarity was computed using the Tanimoto coefficient on the Atomic fingerprints.

**Supplementary Table 4:** The table contains information about the YMDB metabolites predicted to bind at the intracellular cavities IC4 and IC5 of the Ste2 protein. Of note, predictions were made using molecular docking and Gcoupler. It contains information about the YMDB ID, Metabolite name, SMILES (Simplified Molecular Input Line Entry System), and cavity information. Of note, BE refers to binding energies, whereas probabilities were the outcome of Gcoupler.

**Supplementary Table 5:** The table contains information about the list of metabolite-gene pairs from the Kyoto Encyclopedia of Genes and Genomes (KEGG) pathway that is used in this study.

**Supplementary Table 6:** The table contains the results of the untargeted metabolomics. Of note, the values represent normalized and imputed peak intensities of the indicated metabolites.

**Supplementary Table 7:** The table contains the TMM normalized read counts after RNA Sequencing.

**Supplementary Table 8:** The table contains the list of Differentially Expressed Genes (DEGs) of the Control groups, alongside the other standard output from the NOISeq analysis, such as M and D values, mean values of the two conditions, probability ranking, and other gene-associated features.

**Supplementary Table 9:** The table contains the list of Differentially Expressed Genes (DEGs) of the Control vs LST group, alongside the other standard output from the NOISeq analysis, such as M and D values, mean values of the two conditions, probability ranking, and other gene-associated features.

**Supplementary Table 10:** The table contains the list of Differentially Expressed Genes (DEGs) of the LST group, alongside the other standard output from the NOISeq analysis, such as M and D values, mean values of the two conditions, probability ranking, and other gene-associated features.

**Supplementary Table 11:** The table contains information about the comparison of *de novo* drug design and cavity detection tools. Please note, SBDD refers to Structure-Based Drug Design, and LBDD refers to Ligand-Based Drug Design.

**Supplementary Table 12:** The table contains information about the known mutants from the literature that could modulate Ste2 signaling. Of note, the table also contains information about the site of mutation, the nature of the mutation, its position within the intracellular cavities IC4 and IC5, its impact on the Ste2 signaling pathway, and references.

## SUPPLEMENTARY INFORMATION

### Supplementary Information

The supplementary Information provides details of the nucleotide and amino acid Multiple Sequence Alignments (MSAs) of the wild-type *STE2* and the site-directed missense mutants of *STE2*. Each row starts with the sequence identifier followed by the aligned sequence (in chunks) with the ending position of the aligned sequence provided at the end, separated by tabs. The mutation sites are highlighted in red (S75A), yellow (T155D), and blue (L289K). Supplementary Information also contains detailed materials and methods.

